# Selective Regulation of a Defined Subset of Inflammatory and Immunoregulatory Genes by an NF-κB p50-IκBζ Pathway

**DOI:** 10.1101/2024.01.23.576959

**Authors:** Allison E. Daly, George Yeh, Sofia Soltero, Stephen T. Smale

**Affiliations:** Department of Microbiology, Immunology, and Molecular Genetics, Los Angeles, Los Angeles, CA 90095, USA; Molecular Biology Institute, and University of California, Los Angeles, Los Angeles, CA 90095, USA; Howard Hughes Medical Institute, University of California, Los Angeles, Los Angeles, CA 90095, USA

**Keywords:** macrophages, inflammation, transcription, NF-κB, IκBζ

## Abstract

The five NF-κB family members and three nuclear IκB proteins play important biological roles, but the mechanisms by which distinct NF-κB and IκB proteins contribute to selective gene transcription remain poorly understood, especially at a genome-scale level. Using nascent transcript RNA-seq, we observed considerable overlap between p50-dependent and IκBζ-dependent genes in Toll-like receptor 4 (TLR4)-activated macrophages. Key immunoregulatory genes, including *Il6*, *Il1b*, *Nos2*, *Lcn2,* and *Batf,* are among the p50-IκBζ co-dependent genes. IκBζ bound genomic sites occupied by NF-κB dimers at earlier time points. However, p50-IκBζ co-dependence does not coincide with preferential binding of either p50 or IκBζ, as both proteins and RelA co-occupy thousands of genomic sites. A common feature of p50-IκBζ co-dependent genes is a nearby p50/RelA/IκBζ co-bound site exhibiting p50-dependent binding of both RelA and IκBζ. This result and others suggest that IκBζ may act in concert with RelA:p50 heterodimers. Notably, the IκBζ-dependent and p50-IκBζ-co-dependent genes comprise a high percentage of genes that exhibit the greatest differential expression between TLR4-stimulated and tumor necrosis factor receptor (TNFR)-stimulated macrophages. Thus, our genome-centric analysis reveals a defined p50-IκBζ pathway that selectively activates a set of key immunoregulatory genes and serves as an important contributor to the differential TNFR and TLR4 responses.

## INTRODUCTION

Inflammatory responses are activated in diverse mammalian cell types by a wide range of physiological stimuli, including microbial agents, cytokines, and numerous environmental insults. The transcriptional activation of pro-inflammatory genes, which represents a critical component of an inflammatory response, is tailored to the stimulus to defend the host and restore cellular and organismal homeostasis. The tailored transcriptional response is dictated by the sensors of the stimulus, the signaling pathways and transcription factors induced by the sensors, and the poised and stimulus-responsive chromatin state of the genome (Heinz et al. 2015; Glass and Natoli 2016; Monticelli and Natoli 2017; Natoli and Ostuni 2019, Sheu and Hoffmann 2022).

Differential responses to the microbial product, lipid A, and the cytokine, tumor necrosis factor (TNF), have long served as a model for understanding the selectivity of pro-inflammatory gene induction (Covert et al. 2005; Werner et al. 2005). These stimuli, which are sensed by Toll-like receptor 4 (TLR4) and the TNF receptors (TNFRs), respectively, are potent inducers of common transcription factors, including NF-κB, AP-1, and serum response factor (SRF) (Takeuchi and Akira 2010; Hayden and Ghosh 2013; Brenner et al. 2015). However, despite extensive overlap in transcriptional programs, the responses to TLR4 and TNFRs exhibit important differences. Notably, the type 1 interferon (IFN) response and several key cytokines and effector molecules are selectively activated by TLR4 signaling. A prominent contributor to this differential response is TLR4’s selective ability to activate the TRIF signaling pathway and its downstream IRF3 transcription factor (Yamamoto et al. 2002; Fitzgerald et al. 2003; Oshiumi et al. 2003). The TRIF pathway also prolongs the activation of NF-κB complexes, allowing activation kinetics to serve as a major regulator of the selective transcriptional response (Covert et al. 2005; Werner et al. 2005; Cheng et al. 2021).

Although many studies of NF-κB’s role in inflammatory gene transcription have focused on the abundant heterodimer composed of the NF-κB RelA and p50 subunits, the NF-κB family consists of five members – RelA, c-Rel, RelB, p50, and p52 – each of which contains a conserved Rel homology region (RHR) that supports sequence-specific DNA binding and dimerization (Hayden and Ghosh 2013). The five subunits assemble into 15 homodimeric and heterodimeric species, with only the RelA, c-Rel, and RelB subunits containing transactivation domains. Most NF-κB dimers are induced by post-translational mechanisms in response to inflammatory stimuli (Hayden and Ghosh 2013). Phenotypic studies of mice lacking the genes encoding individual NF-κB subunits have revealed distinct immunological defects (Hayden and Ghosh 2013). However, much remains to be learned about dimer-specific functions in vivo and the underlying mechanisms of dimer-specific regulation.

The p50 protein, encoded by the *Nfkb1* gene, like the p52 protein encoded by *Nfkb2*, lacks a transactivation domain. However, it has been proposed to regulate transcription via diverse mechanisms as a subunit of multiple dimeric species. First, the abundant RelA:p50 and c-Rel:p50 heterodimers are thought to support transcriptional activation of numerous pro-inflammatory genes, with the RelA and c-Rel subunits providing the transactivation domain (Hayden and Ghosh 2013). In addition, p50 homodimers have been proposed to serve as transcriptional repressors, due to their absence of a transactivation domain and potential to compete for binding with other NF-κB dimers (Cheng et al. 2011; Zhao et al. 2012). Interactions between p50 homodimers and histone deacetylases have also been suggested to contribute to transcriptional repression (Zhong et al. 2002; Elsharkawy et al. 2010). Finally, p50 has been shown to interact with nuclear inhibitor of NF-κB (IκB) proteins, which serve as co-activators or co-repressors (Hayden and Ghosh 2013).

The mouse and human genomes encode eight IκB proteins, three of which - IκBα, IκBα, and IκBε - play major roles in sequestering NF-κB dimers in the cytoplasm prior to cell stimulation (Hayden and Ghosh 2013). Two other IκB-like proteins are components of the p105 and p100 precursors to the NF-κB p50 and p52 proteins. Of greatest relevance to this study, the remaining three IκB proteins, IκBζ (*Nfkbiz*), IκBNS (IκB8, *Nfkbid*), and Bcl3 (*Bcl3*), are found predominantly in the nucleus and each has been shown to interact with the NF-κB p50 protein (Chiba et al. 2013; Hayden and Ghosh 2013; Schuster et al. 2013; Annemann et al. 2016; Willems et al. 2016). IκBζ primarily serves as a transcriptional co-activator, IκBNS is primarily a co-repressor, and Bcl3 has been proposed to contribute to both activation and repression (Chiba et al. 2013; Hayden and Ghosh 2013; Schuster et al. 2013; Annemann et al. 2016). Notably, other IkBs also enter the nucleus and help regulate NF-κB functions (Huang et al. 2000; Banerjee et al. 2005).

Although much has been learned, the regulatory logic through which different NF-κB dimeric species and nuclear IκB proteins contribute to selective gene regulation remains poorly understood. This knowledge has been difficult to achieve for a variety of reasons, including considerable redundancy between dimeric species, the similarity of binding motifs for many dimers, and the complexity of the mutant phenotypes.

To gain further insight, we began with a genomics approach, in which we performed nascent transcript RNA-seq with bone marrow-derived macrophages (BMDMs) from wild-type (WT) and *Nfkb1^-/-^* mice. This analysis revealed a small number of key inflammatory genes that exhibit strong p50 dependence, including *Il6*, *Il1b*, *Nos2*, *Lcn2*, and *Batf*. Although p50 can contribute to transcription via diverse mechanisms, we found that a surprisingly large fraction of p50-dependent genes also exhibit dependence on IκBζ; other key genes, including *Il12b* and *Il10*, exhibit dependence on only one of the two factors. ChIP-seq analyses reveal extensive co-occupancy of thousands of genomic sites by p50, RelA, and IκBζ. A strong enrichment of p50-IκBζ-co-dependent genes is found near a small subset of genomic sites where both IκBζ and RelA binding are dependent on p50. Together with in vitro interaction and sequential ChIP-seq experiments, these results suggest that IκBζ may interact functionally with p50:RelA heterodimers. In addition, kinetic studies provide genome-scale evidence that IκBζ binds genomic locations that are assembled into open chromatin and pre-bound by NF-κB dimers, with IκBζ binding at functionally important sites correlating with the acquisition of the histone H3K27ac mark representative of the transition to a transcriptionally active state.

Of particular interest, both IκBζ mRNA and the set of p50-IκBζ -co-dependent genes identified in our analyses are among the genes that exhibit the greatest differential expression in response to TLR4 versus TNFR signaling. This finding suggests that poor expression of IκBζ in response to TNFR signaling, partly due to selective instability of IκBζ mRNA (Eto et al. 2003; Yamazaki et al. 2005), represents a third major contributor to differential TLR4 and TNFR responses, in addition to TLR4-selective IRF3 activation and differential NF-κB induction kinetics.

## RESULTS

### NF-κB p50-Dependent Transcription of TLR4-Induced Genes

To uncover the logic through which distinct members of the NF-κB family help coordinate selective transcriptional responses, we first performed RNA-seq with nascent chromatin-associated transcripts isolated from BMDMs prepared from wild-type (WT) and *Nfkb1^-/-^* mice stimulated with lipid A for 0, 0.5, 1, 2, and 6 hrs. We examined nascent transcripts (Bhatt et al. 2012) rather than mRNA to focus on transcriptional differences rather than possible differences in mRNA stability.

In unstimulated BMDMs, the absence of p50 had little effect on nascent transcript levels (Figure 1A). Although p50 homodimers have been proposed to contribute to transcriptional repression (see Introduction), the absence of p50 in unstimulated BMDMs was insufficient to significantly increase or decrease the transcription of any genes. Interestingly, 0.5-hr post-stimulation, only 4 genes exhibit significant (p-adj <0.01) and relatively strong (< 33% relative to WT) dependence on p50 (Figure 1B). *Egr1* stands out among these genes, as it was induced 129-fold in WT BMDMs at the 0.5-hr time point, but only 8-fold in the mutant BMDMs. In contrast, transcript levels for the other three genes, *Hmga1b*, *Slc11a2,* and *Lfng*, were only minimally induced (1.4-fold, *Hmga1b*; 1.7-fold, *Slc11a2*) or repressed (2.5-fold, *Lfng*) at the 0.5-hr time point in WT BMDMs.

**Figure 1.**
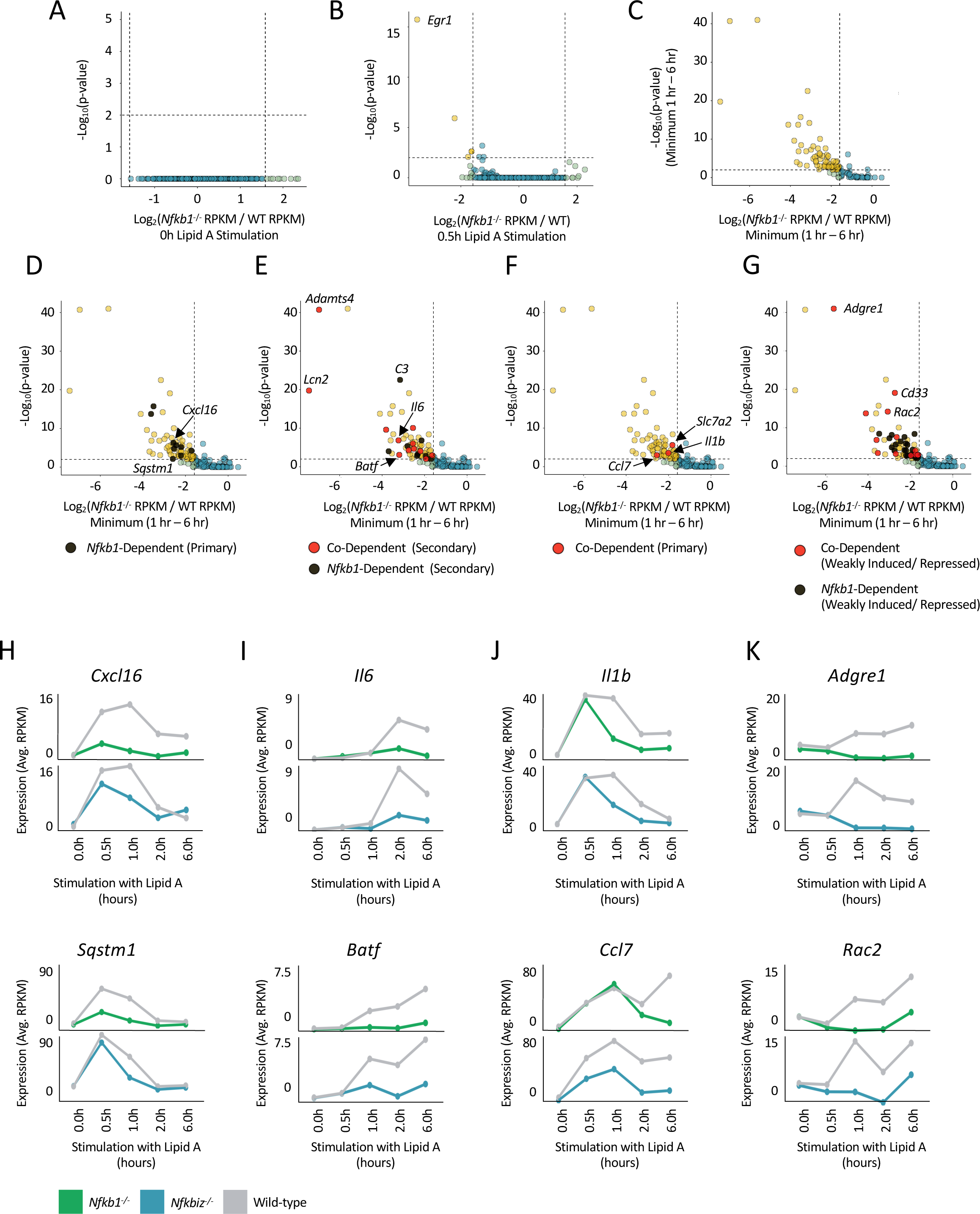
Nascent RNA-seq Analysis of *Nfkb1*^-/-^ and *Nfkbiz*^-/-^ BMDMs. (A) A volcano plot comparing unstimulated *Nfkb1*^-/-^ and WT BMDMs is shown, with dots for all genes considered to be expressed (RPKM >3) in any of the samples analyzed (0, 0.5, 1, 2, or 6 hr). No genes were either positively or negatively regulated to a statistically significant extent prior to stimulation. (B) A volcano plot comparing *Nfkb1*^-/-^ and WT BMDMs stimulated with lipid A for 0.5 hr is shown. Genes with greater dependence and significance than our thresholds (dashed lines) for RPKM ratio (<33%) and p-value (<0.01) are in gold. (C) A volcano plot comparing *Nfkb1*^-/-^ and WT BMDMs stimulated with lipid A for 1, 2, and 6 hr is shown. The values (ratio and p-value) used for each gene were derived from the time point showing the minimum *Nfkb1^-/-^*/WT RPKM ratio (WT RPKM must be >3). Genes with greater dependence and significance than our thresholds (dashed lines) for RPKM ratio (<33%) and p-value (<0.01) are in gold. (D-G) The volcano plot from panel C is shown, with specific gene categories highlighted. (D) *Nfkb1*-dependent primary response genes are highlighted in black. (E) *Nfkb1*-dependent secondary response genes are shown in black and *Nfkb1/Nfkbiz*-co-dependent secondary response genes are in red. (F) *Nfkb1/Nfkbiz*-co-dependent primary genes are highlighted in red. (G) *Nfkb1-*dependent (black) and *Nfkb1/Nfkbiz*-co-dependent (red) genes that are only weakly induced (induction from 1-5-fold) or repressed by lipid A in WT BMDMs are highlighted; these genes do not reach our thresholds for classification as primary or secondary response genes. (H-K) Line graphs are shown displaying nascent transcript kinetics across the induction time-course for representative *Nfkb1*-dependent primary response genes (H), *Nfkb1/Nfkbiz*-co-dependent secondary response genes (I), *Nfkb1/Nfkbiz*-co-dependent primary response genes that display statistical dependence only at late time points after an initial primary response activation (J), and *Nfkb1/Nfkbiz*-co-dependent genes that are only weakly induced or repressed in WT BMDMs (K). Transcript levels are shown for *Nfkb1^-/-^* and *Nfkbiz^-/-^* BMDMs, along with WT BMDMs analyzed in parallel with the mutant cells. Transcript levels represent averages of two biological replicates.

*Egr1* is unlikely to be a p50 target. Instead, its uniquely strong transcript reduction in *Nfkb1^-/-^* BMDMs at the 0.5-hr time point is likely due to its dependence on the mitogen-activated protein kinases (MAPKs), ERK1 and ERK2, which activate serum response factor (SRF)-associated ternary complex factors (TCFs) that are critical for *Egr1* transcription (Hill and Treisman 1995). ERK1/2 activation is dependent on the MAPK kinase kinase, Tpl2, whose activity is strongly deficient in *Nfkb1^-/-^* macrophages. Tpl2 stability requires association with the ankyrin repeat-containing p105 precursor protein encoded by the *Nfkb1* gene (Beinke et al. 2003; Waterfield et al. 2003; Gantke et al. 2011). RNA-seq analyses with lipid A-stimulated *Map3k8^D270A/D270A^* (kinase-inactive Tpl2 mutant) BMDMs (Blair et al. 2022), and with lipid A-stimulated WT BMDMs in the presence of an ERK1/2 inhibitor (Supplemental Figure S1), demonstrated that *Egr1* is by far the most strongly impacted gene in both settings. Thus, *Egr1* deficiency in *Nfkb1^-/-^* BMDMs is likely due to the absence of Tpl2 (note that ERK and the p38 MAPK activate many other TLR4-induced genes in a redundant manner) (Tong et al. 2016).

At 1-, 2-, and 6-hr post-stimulation, only 65 genes (WT RPKM > 3) exhibit strongly diminished transcript levels at one or more time points in the *Nfkb1^-/-^* BMDMs (Figure 1C; < 33% relative to WT; p-value < 0.01). Within this group, the initial induction is strongly diminished for only 6 of the 132 genes we previously identified (Tong et al. 2016) as the most potently induced primary response genes by lipid A in BMDMs, including *Egr1* (the ERK target), *Nfkb2*, *Kdm6b*, *Cxcl16, Sqstm1*, and *Flnb*, as well as for 3 additional genes (*Gadd45b, Tnip1, Gpr132*) with primary response characteristics (Figures 1D, black dots; Supplemental Figure S2). Figure 1H shows representative kinetic profiles for *Cxcl16* and S*qstm1* as examples. The initial induction is also compromised for 6 of the 29 interferon (IFN)-independent secondary response genes we previously defined (i.e. CHX-sensitive induction independent of IFNAR signaling; Tong et al. 2016): *Il6*, *Nos2*, *Il4i1*, *C3*, *Stat5a*, and *Adora2a* (Figures 1E, black and red dots, co-dependence discussed below; Figure 1I, *Il6* example). Strongly diminished induction is also observed with 14 other genes with IFN-independent secondary response characteristics (see Supplemental Figure S2), including *Lcn2*, *Adamts4*, and *Batf* (Figure 1E, black and red dots, co-dependence discussed below; Figure 1I, *Batf* example). None of our previously defined IFNAR-dependent secondary response genes exhibit p50-dependence (data not shown), demonstrating that p50 is not important for a Type I interferon (IFN) response.

Three additional primary response genes, *Il1b*, *Ccl7*, and *Slc7a2* exhibit p50 dependence, but only at later time points (Figure 1F, red dots; Supplemental Figure S2). Transcript profiles for two of these genes, *Il1b* and *Ccl7* (Figure 1J), display their strong, rapid p50-independent induction at the earliest time points examined, followed by p50 dependence at later time points. Thus, although p50 is not required for the initial activation of these genes, their sustained transcription is strongly influenced by p50. The remaining 35 genes that meet our criteria for p50 dependence are induced only weakly or were repressed following lipid A stimulation, with p50 dependence generally observed at the 1-hr time point or later (Figure 1G, 1K). As a general principle, although these genes are induced only weakly, are not induced at all, or are repressed in WT cells following stimulation, their transcript levels at late time points are higher in WT cells than in *Nfkb1^-/-^* cells, demonstrating a role for p50 in their sustained transcription.

To summarize, although p50 is thought to contribute to TLR4-induced transcription via multiple mechanisms, p50-dependent transcription is most pronounced at a small number of inducible genes. Because this group includes several key immunoregulatory genes, including *Il6*, *Il1b*, *Nos2*, *Adamts4*, *Stat5a*, and *Lcn2,* further examination of the underlying mechanism is warranted.

### Initial Analysis of Nuclear IκBs

One mechanism by which p50 might be important for transcription of a limited number of genes is through its reported interactions with the nuclear IκB proteins, which include IκBζ, Bcl3, and IκBNS (Chiba et al. 2013; Hayden and Ghosh 2013; Annemann et al. 2016). To explore a possible role for nuclear IκBs at the p50-dependent genes, we performed nascent transcript RNA-seq with BMDMs from *Nfkbiz^-/-^*(IκBζ), *Bcl3^-/-^*, and *Nfkbid^-/-^*(IκBNS) mice. In all cases, WT and mutant cells were compared in parallel after lipid A stimulation for 0, 0.5, 1, 2, and 6 hr, with two full time-courses performed for each WT and mutant pair. Significant differences are not observed for any genes with either *Bcl3^-/-^* or *Nfkbid^-/-^* BMDMs in this setting (Figure 2A and data not shown). However, in *Nfkbiz^-/-^* BMDMs, 136 genes exhibit strongly diminished transcript levels (WT RPKM > 3; < 33% relative to WT; p-value < 0.01), with 30, 63, and 105 of these genes diminished at the 1, 2, and 6 hr time points, respectively (Figure 2A, 2B). A subset of these genes were previously shown to be IκBζ-dependent in macrophages (Yamamoto et al. 2004; Chiba et al. 2013; Annemann et al. 2016). No genes were diminished at the 0 and 0.5 hr time points, consistent with prior knowledge that *Nfkbiz* is only weakly expressed in unstimulated BMDMs and is a potently induced primary response gene (Tong et al. 2016).

**Figure 2.**
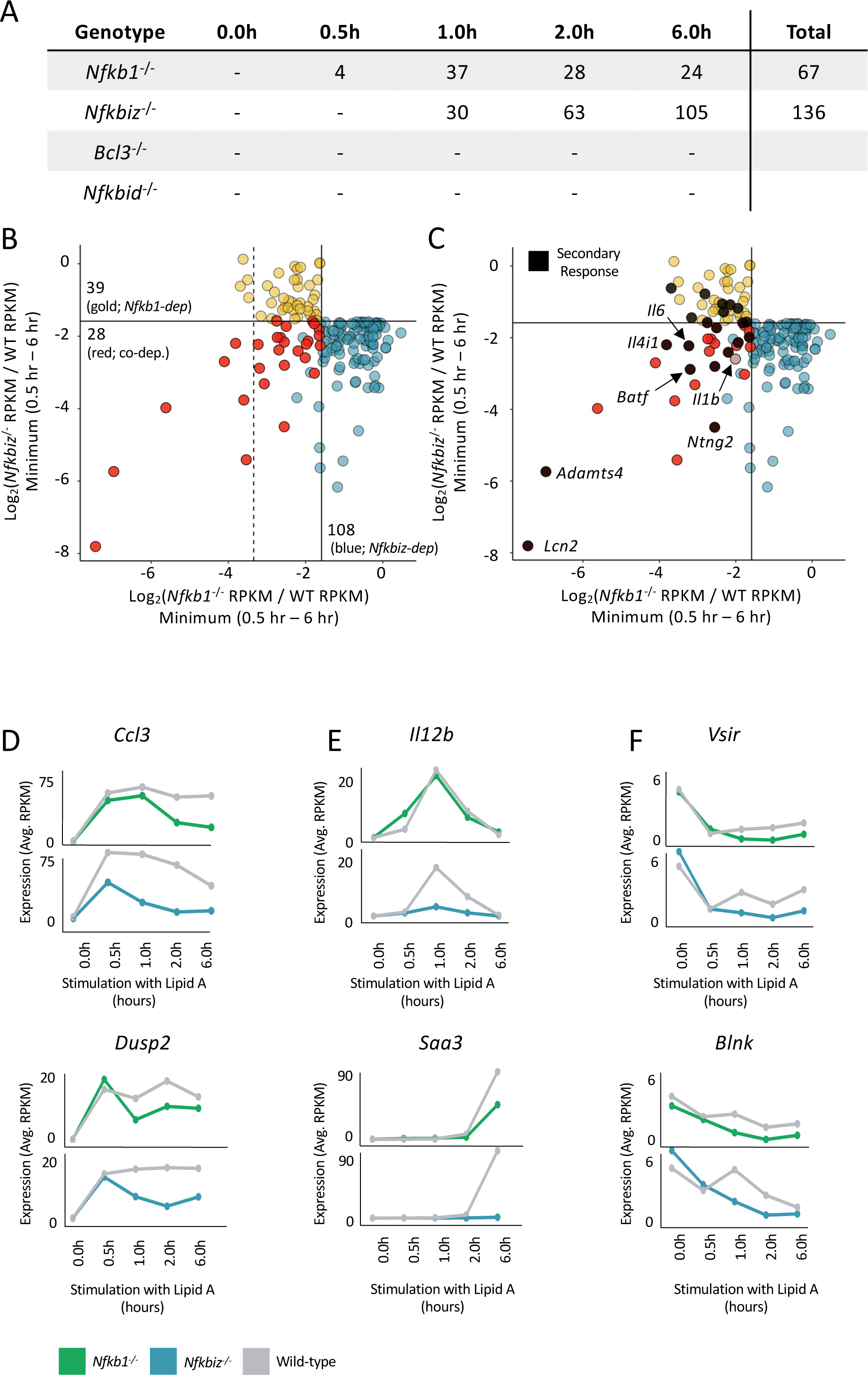
Nascent RNA-seq Analysis of *Nfkbiz*^-/-^, *Bcl3^-/-^*, and *Nfkbid^-/-^* BMDMs. (A) The total number of *Nfkb1*^-/-^, *Nfkbiz*^-/-^, *Bcl3*^-/-^, or *Nfkbid*^-/-^ dependent genes (< 33% relative to WT; p-value <0.01; RPKM >3) at each lipid A stimulation time point is shown. (B) The scatter plot reveals genes exhibiting *Nfkb1* and/or *Nfkbiz* co-dependence. Log2 ratios of *Nfkb1*^-/-^ RPKM/WT RPKM (x-axis) and *Nfkbiz*^-/-^ RPKM/WT RPKM (y-axis) were plotted. The values used for each gene were derived from the time point (0.5, 1, 2, or 6 hr) resulting in the smallest RPKM ratio (WT RPKM must be >3). Solid lines represent an RPKM ratio of 0.33. Genes showing *Nfkb1/Nfkbiz* co-dependence, *Nfkb1* dependence, or *Nfkbiz* dependence are in red, gold, and blue, respectively. To be considered dependent or co-dependent, a gene must be below both our ratio (0.33) and p-value (0.01) thresholds. Thus, genes colored blue in the lower left quadrant are not colored red because their p-values are not below the p-value threshold for *Nfkb1* dependence. The dashed line corresponds to an *Nfkb1*^-/-^ RPKM/WT RPKM ratio of 0.1, to highlight the fact that 7 of 11 genes with the greatest *Nfkb1* dependence display *Nfkb1/Nfkbiz* co-dependence. (C) The scatter plot from panel B is reproduced with *Nfkb1*-dependent secondary response genes in black. *Il1b*, a primary response gene with *Nfkb1/Nfkbiz* co-dependence at late time points is also noted. (D-F) Line graphs are shown displaying nascent transcript kinetics across the induction time-course for representative *Nfkbiz*-dependent primary response genes (D), *Nfkbiz*-dependent secondary response genes (E), and genes repressed by lipid A in WT BMDMs that display statistical *Nfkb1*/*Nfkbiz*-co-dependence. Transcript levels are shown for *Nfkb1^-/-^* and *Nfkbiz^-/-^* BMDMs, along with WT BMDMs analyzed in parallel with the mutant cells. Transcript levels represent averages of two biological replicates.

Interestingly, 28 of the 67 genes (42%) exhibiting strong p50 dependence between 0.5 and 6 hr post-stimulation are also dependent on IκBζ (Figure 2B, red dots, Supplemental Figure S2). In fact, p50/IκBζ co-dependence is observed at 7 of the 11 genes (64%) exhibiting the strongest dependence on p50 (*Nfkb1^-/-^* vs. WT RPKM ratio <10%; see vertical dashed line in Figure 2B, Supplemental Figure S2). Most notably, 12 of the 20 p50-dependent secondary response genes (60%) exhibit IκBζ co-dependence (Figure 2C; black dots; Supplemental Figure S2), including *Il6*, *Il4i1*, *Adamts4*, *Nos2*, *Batf*, *Lcn2*, and *Slc11a2* (see representative kinetic profiles in Figure 1I). IκBζ did not contribute to the initial induction of any of the p50-dependent primary response genes that exhibited p50 dependence during the primary response (Figure S2; representative profiles in Figure 1H), consistent with the fact that the *Nfkbiz* gene is itself a primary response gene. However, all 3 primary response genes that exhibited delayed p50 dependence, *Il1b*, *Ccl7*, and *Slc7a2*, exhibited IκBζ co-dependence (Figure 1F, 2C; Supplemental Figure S2; representative profiles in Figure 1J).

In addition to the genes that show clear co-dependence on both p50 and IκBζ for their activation, 18 other primary response genes, 18 secondary response genes, 47 weakly induced genes, and 25 genes repressed upon lipid A stimulation exhibit IκBζ dependence (based on our dependence criteria) but without p50 co-dependence (Figures 2B, blue dots; Supplemental Figure S3). All of the primary response genes that exhibit strong dependence only on IκBζ, including *Ccl2*, *Ccl3*, Ccl7, *Dusp1*, and *Dusp2*, exhibit IκBζ dependence almost exclusively at late times after their initial primary response activation (Figure 2D; Supplemental Figure S3). As shown in Figure 2D for representative examples, transcript levels for some of these genes are also reduced in *Nfkb1^-/-^* cells, but by a smaller magnitude that does not meet our dependence criteria.

Secondary response genes dependent only on IκBζ included two prominent cytokine genes, *Il12b*, *Il10*, and the serum amyloid gene, *Saa3* (Supplemental Figure S3; examples in Figure 2E). The weakly induced and repressed genes that exhibit IκBζ dependence generally exhibit dependence for their sustained transcription at late time points (Supplemental Figure S3; examples in Figure 2F). Notably, several of the induced genes exhibiting p50 and/or IκBζ dependence exhibit both primary and secondary response characteristics (i.e. early CHX-independent induction followed by later CHX-dependent super-induction; data not shown).

To summarize, p50/IκBζ co-dependence is observed at 42% of p50-dependent genes, including 64% of the genes exhibiting the strongest p50 dependence. Other genes reach our dependence criteria for only one of the two factors. For secondary response genes that exhibit co-dependence, both factors are needed for the initial transcriptional activation. In contrast, primary response genes that exhibit co-dependence, including *Il1b*, require both factors only for sustained transcription after initial primary response activation. Although co-dependence is observed for only a minority of genes that exhibit dependence on either factor, one could argue that the prevalence of co-dependence is surprising, given the diverse mechanisms by which p50 is thought to contribute to gene transcription. An interest in better understanding the close collaboration between p50 and IκBζ is heightened by the frequent mutation of, and association with, the *NFKBIZ* gene in human disease, including cancer and ulcerative colitis (Kakiuchi et al. 2020; Nanki et al. 2020; Rheinbay et al. 2020)

### Analysis of *Nfkb2^-/-^* and *Nfkb1^-/-^Nfkb2^-/-^* Macrophages

Many NF-κB target genes activated by p50-containing dimers may not exhibit p50-dependent transcription in lipid A-stimulated BMDMs due to redundancy with p50’s closely related paralog, p52. Redundancy with p52 may also contribute to the existence of a set of IκBζ-dependent/p50-independent genes. To gain insight into the contribution of p52, we first performed RNA-seq with *Nfkb2^-/-^* BMDMs. The results revealed a small number of p52-dependent genes. However, none of these genes exhibited IκBζ dependence (data not shown). This result demonstrates the absence of genes that are uniquely p52/IκBζ co-dependent.

We were unable to successfully breed *Nfkb1^-/-^Nfkb2^-/-^*mice, consistent with prior evidence of poor viability (Franzoso et al. 1997A). Therefore, to examine possible redundancy between p50 and p52, we created a J2 retrovirus-immortalized macrophage line from *Nfkb1^-/-^*mice and then used CRISPR-Cas9 editing to disrupt the *Nfkb2* gene in this line (Supplemental Figure S4). RNA-seq analysis (mRNA) comparing the *Nfkb1^-/-^Nfkb2^-/-^*line and the parental *Nfkb1^-/-^* line to a WT J2-transformed macrophage line revealed that the combined p50 and p52 deficiency yields reduced transcript levels at a larger number of inducible genes than observed in the absence of p50 alone (Supplemental Figure S4D), confirming the existence of significant redundancy between the two paralogs. However, about 70% of strongly induced genes remain largely unaffected (Supplemental Figure S4), either because they can be activated by NF-κB dimers that do not include either p50 or p52 or because their induction is entirely NF-κB independent.

Focusing on IκBζ -dependent genes, we found that only a few additional IκBζ-dependent/p50-independent genes exhibit reduced transcript levels in the *Nfkb1^-/-^Nfkb2^-/-^* line (Supplemental Figure S4D). This result demonstrates that the p50-independence of IκBζ-dependent/p50-independent genes is not due to redundancy between p50 and p52.

### Initial p50, IκBζ, and RelA Chip-Seq Analyses

To help elucidate the mechanisms underlying the regulation of p50/IκBζ co-dependent genes, we performed ChIP-seq with p50, RelA, and IκBζ antibodies in BMDMs stimulated with lipid A for 0, 0.5, 1, and 2 hr. We observed 2,311, 6,189, and 3,310 reproducible peaks (peak score >19 and RPKM >3 in 2/2 biological replicates at one or more time points) with p50, RelA, and IκBζ antibodies, respectively (Figure 3A, right). The different peak numbers may be due to a combination of different antibody qualities or different numbers of genomic interactions. Despite the different peak numbers, extensive overlap of genomic interactions is observed for the three proteins (Figure 3B). Importantly, the large number of peaks and their extensive overlap demonstrate that p50/IκBζ-co-dependent and IκBζ-dependent transcription is not simply due to highly selective genomic interactions.

**Figure 3.**
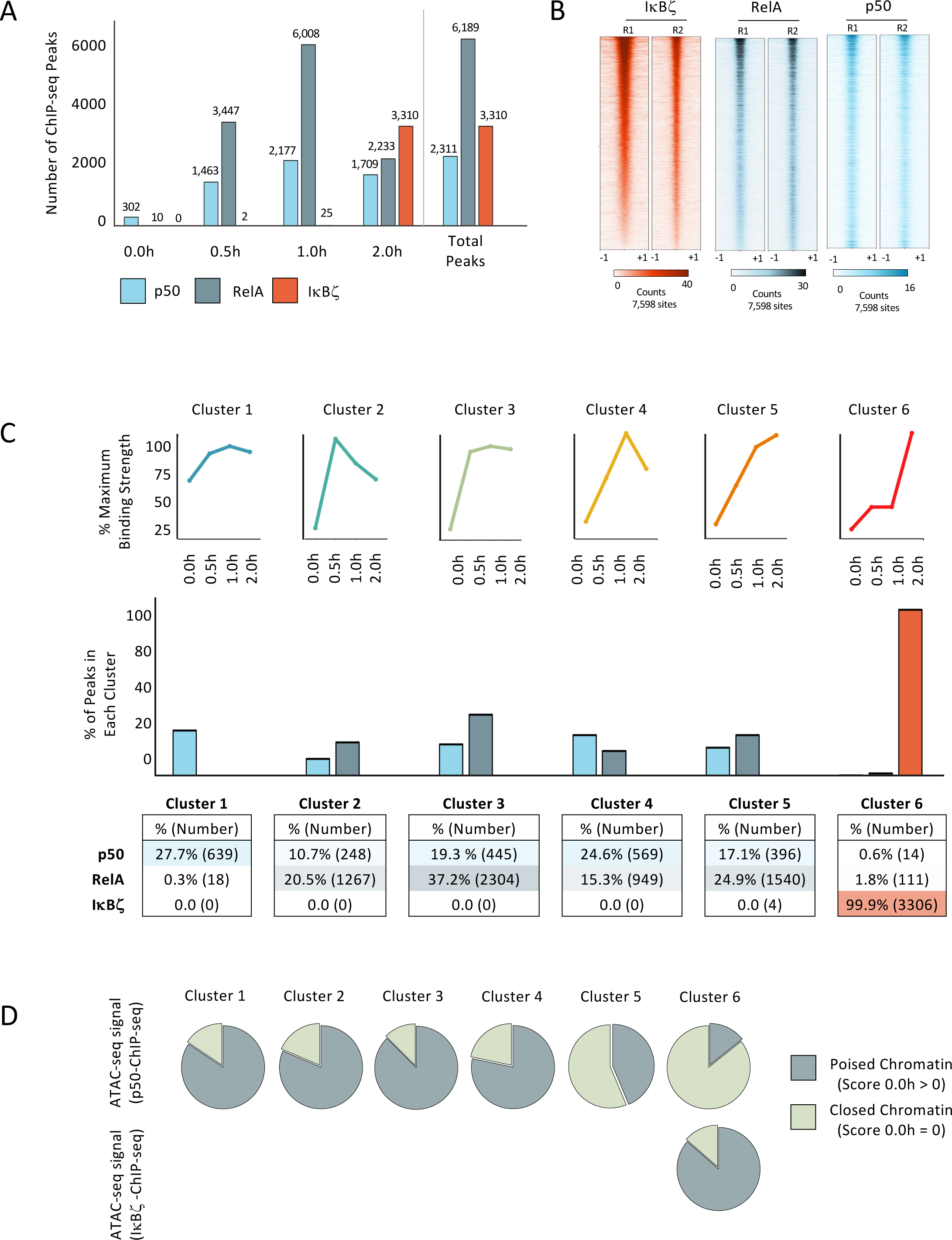
RelA, p50, and IκBζ ChIP-seq Analysis. (A) The number of reproducible binding sites (PS > 19 and RPKM > 3 in 2/2 replicates) at each time point (0, 0.5, 1, and 2 hr) is plotted. The total number of unique binding sites for each protein is shown (right). (B) Heatmaps are shown from two biological replicates of ChIP-seq experiments for IκBζ, RelA, and p50 in cells stimulated for 2 hr. The heatmaps contain 7,598 peaks, which includes all unique peaks bound by RelA, p50, or IκBζ. All heatmaps are ranked in descending order based on the binding strength in IκBζ replicate 1 (R1). (C) IκBζ, RelA, and p50 ChIP-seq peaks (average RPKM of two replicates for each protein and each time point) were combined and examined by k-means cluster analysis to identify six kinetic clusters. Line graphs are shown to represent the six clusters. The percentages of ChIP-seq peaks for p50, RelA, or IκBζ found in each of the six kinetic clusters from panel C are shown in bar graphs. Percentages and numbers are shown at the bottom. (E) At the top, pie graphs show the fraction of p50 ChIP-seq peaks in each of the six kinetic clusters found at genomic regions that either exhibit chromatin accessibility or inaccessibility by ATAC-seq (i.e. called ATAC-seq peaks; peak score >0 or =0, respectively) in unstimulated BMDMs. Most p50 ChIP-seq peaks in kinetic clusters 1-4 exhibit ATAC-seq sensitivity in unstimulated cells (i.e. poised chromatin). In contrast, greater than half of p50 ChIP-seq peaks in clusters 5 and 6 occur at locations that are inaccessible in unstimulated cells. The pie graph at the bottom shows that, although IκBζ binding always aligns with kinetic cluster 6, locations bound by IκBζ are usually accessible in unstimulated cells (i.e. poised chromatin).

To better understand the relationships between the three proteins, we compared peak numbers at each time point (Figure 3A, left). In unstimulated cells, 302 p50 peaks are observed, consistent with evidence that p50 homodimers but not other NF-κB dimeric species localize to the nucleus prior to stimulation (Kang et al. 1992; Ten et al. 1992; Zhong et al. 2002). At 0.5 and 1 hr post-stimulation, increasing numbers of p50 and RelA peaks are observed, consistent with rapid nuclear translocation of diverse NF-κB dimeric species. In contrast, IκBζ peaks are largely restricted to the 2-hr time point, consistent with the time needed to express IκBζ protein from the lipid A-induced *Nfkbiz* gene (Figure 3A).

### IκBζ Binds Genomic Sites Bound Earlier by NF-κB

Next, k-means cluster analysis with all p50, RelA, and IκBζ binding sites throughout the time-course allowed the peaks observed for each protein to be classified into six distinct kinetic clusters (Figure 3C, top). Cluster 1, representative of peaks first observed in unstimulated cells, is apparent only at a subset (639, 27.7%) of the p50 peak profiles (Figure 3C, middle and bottom), consistent with the fact that only p50 is present in the nucleus prior to stimulation. Substantial percentages of both the p50 and RelA peak profiles are highly prevalent in clusters 2-5, in which strong binding is first observed at either the 0.5 or 1-hr time point (Figure 3C). In contrast, IκBζ peaks are restricted almost entirely (99.9%) to cluster 6, in which binding is low or absent prior to the 2-hr time point (Figure 3C). Notably, fewer than 2% of p50 and RelA peaks aligned with this kinetic profile (Figure 3C). Thus, given that a very high percentage of IκBζ ChIP-seq peaks overlap with p50 and/or RelA peaks, the results suggest that IκBζ typically binds genomic sites bound at earlier time points by NF-κB dimers (see below).

Next, we used ATAC-seq results to examine the chromatin state of genomic sites bound by p50, RelA, and IκBζ. Most p50 peaks within clusters 1-4 are found at genomic regions with ATAC-seq peaks in unstimulated cells (Figure 3D), consistent with extensive evidence that NF-κB dimers often bind regulatory regions assembled into poised chromatin (Ghisletti et al. 2010; Heinz et al. 2010). In contrast, p50 peaks within clusters 5 and 6, which exhibit relatively late p50 binding, are found at genomic regions that lack ATAC-seq peaks in unstimulated cells (Figure 3D). This finding suggests that the delayed p50 binding at peaks within these clusters is associated with a frequent need for nucleosome remodeling prior to NF-κB binding. Importantly, although IκBζ peaks align almost exclusively with cluster 6, these peaks are found at genomic regions that exhibit ATAC-seq accessibility representative of open chromatin in unstimulated cells (Figure 3D, Cluster 6, bottom). This finding further strengthens the notion that IκBζ typically binds regions that are already assembled into open chromatin and already associated with an NF-κB dimer. These findings from genomic studies are consistent with a prior study performed with representative genes, which revealed that IκBζ acts after nucleosome remodeling (Kayama et al. 2008), but inconsistent with another prior study that suggested a role for IκBζ in nucleosome remodeling (Tartey et al. 2014).

### Analysis of Preferential Genomic Interactions by p50 and RelA

The above results suggest that p50, RelA, and IκBζ bind together at thousands of genomic locations, with an NF-κB dimer binding first, followed by association of IκBζ after transcriptional induction of the *Nfkbiz* gene. To further elucidate the characteristics of genomic regions bound by these proteins, we asked whether IκBζ binding correlates with preferential binding by p50 or RelA. For this analysis, we first selected the strongest reproducible 2,311 peaks from the p50 and RelA ChIP-seq experiments, which generated 3,541 total peaks when combined. We then calculated the p50/RelA RPKM ratio at each genomic region containing a called peak for either protein (Figure 4A).

**Figure 4.**
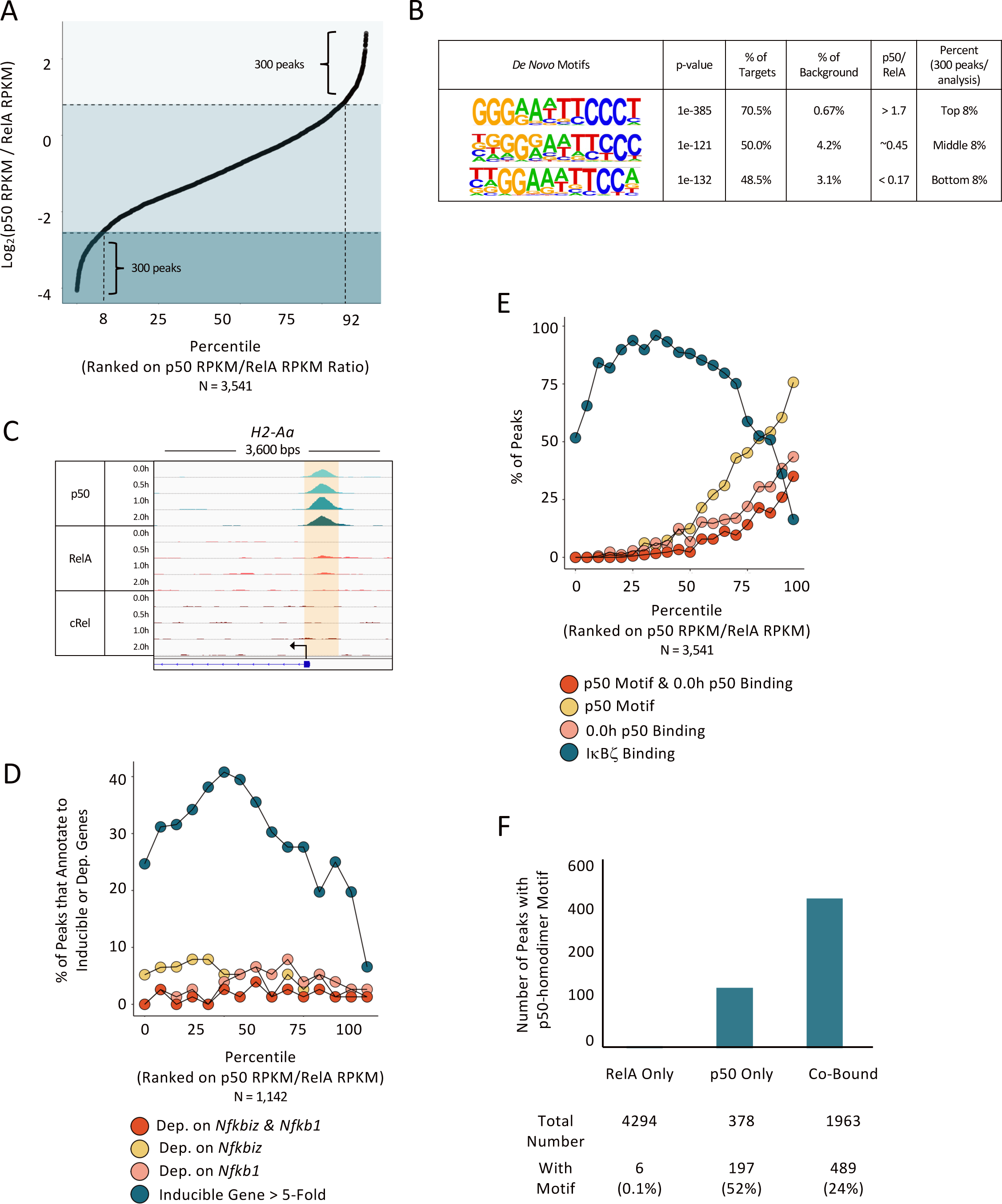
Relationship between Preferential p50 or RelA Binding and Transcriptional Dependence. (A) All reproducible p50 binding sites (PS > 19 and RPKM > 3 in 2/3 replicates; 2,311 peaks) and the top 2,311 reproducible RelA binding sites were combined (total of 3,541 peaks) and ordered based on the 1-hr average p50 RPKM/RelA RPKM ratio. The 300 peaks (8% of total) with the strongest preference for either p50 (top right) or RelA (bottom left) are indicated. (B) De novo motif analyses were performed with genomic sequences underlying the 300 peaks with the strongest p50 preference (top 8%), the strongest RelA preference (bottom 8%), and with 300 peaks showing no preference (middle 8%). (C) Browser tracks at the *H2-Aa* locus are shown as an example of p50 preferential binding, with strong p50 peaks at the *H2-Aa* promoter in unstimulated and stimulated BMDMs, and no peaks for RelA or c-Rel. (D) Each ChIP-seq peak from panel A was annotated to its closest gene. Peaks located within 5 kb of the TSS for a gene (1,142 peaks) were included in this analysis and the remaining peaks were excluded. The included peaks were divided into 15 equal bins. The percentage of peaks in each bin that annotates to a gene induced by lipid A by >5-fold are shown in blue, revealing that peaks preferentially bound by p50 infrequently annotate to inducible genes. The percentages of peaks in each bin that annotate to *Nfkb1/Nfkbiz*-co-dependent (red), *Nfkbiz*-dependent (gold), and *Nfkb1*-dependent (pink) genes are also shown, revealing no correlation with preferential binding. (E) p50 and RelA ChIP-seq peaks from panel A were divided into 20 equal bins. The percentage of peaks in each bin that coincide with an IκBζ ChIP-seq peak are shown in blue, demonstrating that IκBζ binds infrequently to regions displaying p50-preferential binding. The percentages of peaks in each bin that exhibit p50 ChIP-seq peaks in unstimulated cells is shown in pink. The percentage of peaks in each bin that contain the motif that most strongly supports p50 homodimer binding (three G/C bps in each half-site separated by five bps) is shown in gold, and the percentage that contains both a p50 homodimer motif and displays p50 binding in unstimulated cells is in red. (F) The number of genomic locations that exhibit reproducible peaks (PS >19 and RPKM >3 in 2/3 samples) for only RelA (4294), for only p50 (378), and for both p50 and RelA (1963) are shown, along with the number of peaks in each group that contain a p50 homodimer motif (three GC bps in each half-site separated by five bps). This analysis shows that, although the probability of finding a p50 homodimer motif at a location that displays only a p50 ChIP-seq peak is the highest (52% of these peaks), a larger number of locations with both p50 and RelA ChIP-seq peaks contain p50 homodimer motifs (489) than at locations with only a p50 ChIP-seq peak (197).

Motif enrichment analysis was performed with the 300 peaks representing either the largest or smallest p50/RelA RPKM ratios, as well as the 300 peaks in the middle of the ratio spectrum (Figure 4B). The bin with the greatest p50-preferential binding (largest p50/RelA ratio, Figure 4B, top) is enriched in NF-κB motifs representative of p50 homodimer binding, with three G:C-bps in each half-site (structural and biochemical studies revealed that p50 subunits strongly associate with three G:C bps within a half-site, whereas RelA associates with only two G:C bps [Huxford and Ghosh 2009]). The bin with the greatest RelA-preferential binding (smallest p50/RelA ratio, Figure 4B, bottom) is enriched in motifs representative of RelA homodimer binding, with two G:C-bps in each half-site. A bin with comparable binding by p50 and RelA (Figure 4B, middle) is enriched in motifs that less clearly represent either p50 or RelA interactions. Browser tracks displaying an example of a p50-preferential binding site at the *H2-Aa* promoter are shown in Figure 4C.

We next divided the ChIP-seq peaks examined in Figure 4A into 15 equal bins and calculated the percentage of peaks within each bin that annotate to a TLR4-induced gene (fold induction >5) using a nearest gene approach (Figure 4D). Interestingly, ChIP-seq peaks that exhibit the greatest p50-preferential binding (Figure 4D, right) are the least likely to annotate to induced genes, suggesting that p50 homodimer interactions do not typically contribute to TLR4-induced transcription.

We also observed that p50-preferential binding correlates with an increased probability of p50 binding in unstimulated cells. Specifically, in the bin with peaks displaying the greatest p50 preferential binding, 43% of peaks are at genomic locations exhibiting p50 peaks in unstimulated cells (Figure 4E, pink). Furthermore, in this bin, 76% of the peaks coincide with a DNA motif expected to support p50 homodimer binding (Figure 4E, yellow), with successively smaller percentages in bins with lower p50/RelA binding ratios. Overall, 35% of peaks in this bin both contain a homodimer motif and exhibited a p50 peak in unstimulated cells (Figure 4E, red).

The above results reveal that p50 homodimer motifs are prevalent at sites displaying p50-preferential binding in stimulated cells. However, although the probability of finding a p50 homodimer motif is highest in bins displaying p50-preferential binding, this represents a relatively small number of p50-binding sites with p50 homodimer motifs. Specifically, 197 of 378 (52%) of the p50-called peaks that lack RelA-called peaks contain a p50 homodimer motif (Figure 4F). In comparison, 489 of 1963 (24%) of the p50 called peaks that also display a RelA called peak contain a p50 homodimer motif. Thus, a p50 homodimer motif does not strongly predict p50-preferential binding. One possibility is that, following cell stimulation, the abundance of RelA:p50 heterodimers greatly exceeds the abundance of p50 homodimers, leading to RelA:p50 heterodimer binding to homodimer motifs.

To summarize, the results in Figure 4 demonstrate that p50-preferential and RelA preferential binding can be observed in lipid A-stimulated macrophages, with bins containing preferentially bound sites enriched in motifs predicted to represent binding of p50 homodimers or RelA homodimers, respectively. p50-preferential peaks are least likely to be near a lipid A-induced gene, but they are the most likely to correlate with p50 binding in unstimulated cells. Moreover, the presence of a p50 homodimer motif does not predict the presence of p50-preferential binding.

### IκBζ Genomic Interactions and IκBζ-Dependent Genes are Enriched at Regions that Support p50 and RelA Co-Binding

The above insights provide a framework for evaluating the landscape of IκBζ binding. Toward this goal, we next explored the prevalence of IκBζ ChIP-seq peaks across the RelA-p50 binding ratio profile. This analysis reveals that IκBζ binding is enriched at sites that support binding by both p50 and RelA (p50/RelA ratio of 0.3 – 1.6) and is observed much less frequently in bins with either a strong p50 or a strong RelA preference (Figure 4E). This result suggests the possibility that IκBζ may not preferentially bind p50 homodimers.

To examine the relationship between preferential p50-RelA binding and transcription, we returned to the gene dependence classifications from Figure 2B and asked whether p50/IκBζ-co-dependent, IκBζ-dependent, and p50-dependent genes correlate either with genomic regions that exhibit preferential p50 binding. For this analysis, each peak was annotated to its closest gene (but only if a gene’s TSS is located within 5 kb of the ChIP-seq peak, to increase confidence in the assignments), and the prevalence of peaks annotating to p50/ κBζ-, IκBζ-, and p50-dependent genes was analyzed across the spectrum of p50-RelA RPKM ratios. This analysis reveals that peaks near all three gene classes are broadly distributed, with no preference toward peaks preferentially bound by either RelA or p50 (Figure 4D). Together, the distribution of IκBζ binding and the distribution of peaks near genes that exhibit dependence on p50 and/or IκBζ suggest that selective regulation by these factors may not rely on p50 homodimers.

### p50- and IκBζ-Dependent Genes Correlate with p50-Dependent IκBζ Genomic Interactions

The above analyses provide important insight into the genomic landscape of p50, RelA, and IκBζ binding, but failed to reveal the mechanism by which a key group of inflammatory genes exhibit transcriptional dependence on p50 and/or IκBζ. We therefore extended our analysis by examining the p50 dependence of IκBζ genomic interactions. For this analysis, IκBζ ChIP-seq was performed in parallel with WT and *Nfkb1^-/-^* BMDMs. Surprisingly, only about 5% of IκBζ genomic interactions (187 peaks) exhibited strong dependence (< 33% binding of IκBζ in *Nfkb1*^-/-^ cells relative to WT cells) on p50, with the remaining IκBζ peaks exhibiting either weak or no dependence (Figure 5A).

**Figure 5.**
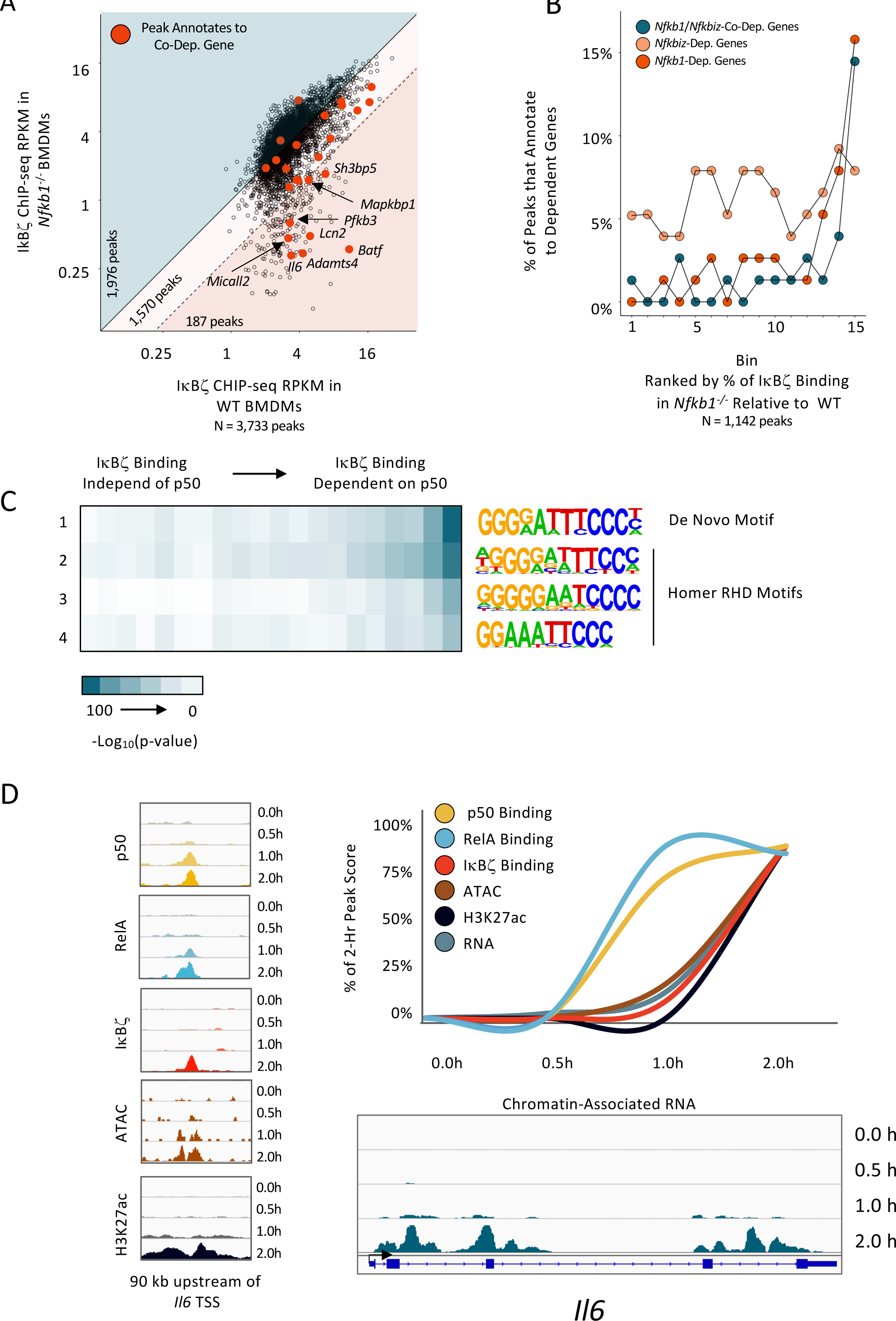
p50 Dependence of IκBζ Binding and Time Course of Events. For all IκBζ ChIP-seq peaks (PS > 19 and RPKM >3 in 2/3 replicates) in WT macrophages stimulated with lipid A for 2.0h, the average RPKM in WT (x-axis) versus *Nfkb1*^-/-^ (y-axis) macrophages is plotted. The blue background highlights 1,976 peaks with >100% IκBζ RPKM in *Nfkb1*^-/-^ versus WT macrophages. The light pink background highlights 1,570 IκBζ binding sites with a mild dependence on *Nfkb1* (33-100% relative to WT). The pink background highlights 187 IκBζ peaks (5% of total) with a strong dependence on p50 (< 33% binding strength relative to WT). Dots colored red indicate binding sites that annotate to *Nfkb1*/*Nfkbiz*-co-dependent genes. Although multiple peaks may annotate to a single gene, only one peak per gene is highlighted in red: the peak with the maximum change in IκBζ binding between *Nfkb1*^-/-^ and WT BMDMs. (A) Quantification of the results in panel A is shown. IκBζ peaks (PS >19 and RPKM >3 in 2/3 replicates; restricted to peaks within 30 kb of a gene expressed at >3 RPKM, with only one peak included per gene based on greatest *Nfkb1-*dependence of IκBζ binding) were placed in 15 bins based on relative dependence on *Nfkb1*. In each bin, the percentage of peaks that annotate to an *Nfkb1*/*Nfkbiz*-co-dependent (blue), *Nfkbiz-*dependent (gold), and *Nfkb1*-dependent (red) gene is plotted. (C) Motif analysis is shown for IκBζ peaks placed into 20 bins based on relative dependence on p50. Three NF-κB motifs from HOMER and one de novo motif were used in this analysis. The heatmap corresponds to the -Log_10_(p-value) for motif enrichment. (D) Properties of a representative p50-dependent IκBζ binding site located 90 kb upstream of the *Nfkb1*/*Nfkbiz*-co-dependent *Il6* gene are displayed. At the left, genome browser tracks show ChIP-seq tracks for p50, RelA, IκBζ, and H3K27Ac, and ATAC-seq tracks, at 0, 0.5, 1, and 2 hr after lipid A stimulation. The ChIP-seq, ATAC-seq and chromatin-associated transcript RNA-seq data are quantified in the line graph using peak scores for ChIP-seq and ATAC-seq data and RPKM for RNA-seq data. Genome browser tracks showing chromatin-associated transcript data for the *Il6* gene at each time point are shown at the bottom.

To evaluate the possible significance of this small set of p50-dependent IκBζ binding events, we examined their locations in relation to p50/IκBζ-, IκBζ-, and p50-dependent genes. Interestingly, IκBζ ChIP-seq peaks near p50-IκBζ co-dependent and p50-dependent genes are frequently among the small fraction of IκBζ ChIP-seq peaks that require p50 (Figure 5A; peaks near co-dependent genes in red). To quantify this finding, IκBζ ChIP-seq peaks from this experiment, located within 30 kb of an expressed gene, were separated into 15 equal bins on the basis of their quantitative dependence on p50. The percentage of peaks within each bin that annotate to genes with transcriptional p50/IκBζ co-dependence, p50-dependence, and IκBζ-dependence was then calculated. In the bin containing IκBζ peaks that exhibit the strongest requirement for p50 (bin 15), 14% of the peaks annotate to genes with transcriptional p50/IκBζ co-dependence, 16% annotate to genes with transcriptional p50-dependence, and 8% annotate to genes with transcriptional IκBζ-dependence (Figure 5B). Thus, in total, a remarkable 38% of peaks in the IκBζ peak bin exhibiting the strongest binding dependence on p50 annotate to a gene exhibiting transcriptional dependence on p50 and/or IκBζ. For transcriptional p50/IκBζ co-dependence and p50-dependence, this represents a large enrichment, as fewer than 3% of IκBζ peaks in bins 1-12 annotate to either p50/IκBζ co-dependent or p50-dependent genes. These results demonstrate that, although IκBζ genomic interactions can be observed by ChIP-seq at thousands of genomic locations, we can enrich for those that support p50/IκBζ-co-dependent or p50-dependent transcription by examining the dependence of IκBζ binding on p50.

The above results suggest that p50-dependence of IκBζ binding may be predictive of a specific interaction or a binding conformation that supports IκBζ function (see Discussion). To explore these possibilities, we examined binding strength, binding preferences, and binding motif differences between p50-dependent and p50-independent IκBζ genomic interaction sites. First, IκBζ, RelA, and p50 peak score distributions were comparable at p50-dependent and p50-independent IκBζ binding sites (data not shown). In addition, no difference in p50 or RelA preferential binding was observed at p50-dependent versus p50-independent IκBζ binding sites (data not shown).

To compare motif enrichment at p50-dependent versus p50-independent IκBζ binding sites, we separated all IκBζ binding sites into 20 bins based on their magnitude of p50 dependence and then examined the enrichment of four NF-κB motifs in each bin (three NF-κB motifs from the Homer Motif Software and a de novo motif generated from our data). We found the strongest enrichment of all four motifs in bins containing IκBζ peaks with the greatest p50-dependence (Figure 5C). This result suggests that functional interactions between IκBζ and NF-κB occur at NF-κB consensus motifs. The relatively low enrichment of NF-κB motifs at p50-independent IκBζ peaks raises the possibility that IκBζ binding to these sites often does not require an interaction with an NF-κB dimer. However, NF-κB co-occupies a high percentage of these sites with IκBζ (data not shown), suggesting instead that both IκBζ and an NF-κB dimer occupy these sites, often in the absence of an NF-κB consensus motif.

Finally, we explored the relationship between the dependence of IκBζ binding on p50 dependence and H3K37ac, a histone modification frequently used as a marker of a transcriptionally active state. At a genome-wide scale, the kinetics of IκBζ binding does not correlate with the kinetics of the histone H3K27ac signal. However, if the vast majority of IκBζ interactions are not functionally important, this observation may not be meaningful. If we instead focus on the small number of IκBζ peaks that annotate to p50 IκBζ co-dependent genes, the kinetics of IκBζ binding correlates much more closely with the kinetics of the H3K27ac modification. A putative *Il6* enhancer that displays strong p50-dependent IκBζ binding provides an example of this and other kinetic relationships (Figure 5D). At this enhancer, p50 and RelA peaks are both readily observed at the 1-hr time point, but IκBζ binding, H3K27ac, and nascent transcripts are barely detectable until the 2-hr time point. The ATAC-seq peak is not called until the 2-hr time point, but an examination of the ATAC-seq tracks (Figure 1D, left) shows that accessibility is substantially elevated by the 1-hr time point, suggesting that initial chromatin opening coincides with RelA and p50 binding. Together, these results suggest that, at IκBζ binding sites that are likely to be functionally important, NF-κB-dimer binding and chromatin opening precede, IκBζ binding, H3K27ac, and transcription. Thus, a kinetic relationship between IκBζ binding and the H3K27ac modification may be another feature of functionally important IκBζ binding events, along with the p50-dependence of IκBζ binding.

### In Vivo and In Vitro Analysis of IκBζ Interactions with p50 and RelA

The results in Figure 3 demonstrate that IκBζ co-occupies thousands of genomic sites with both the p50 and RelA subunits of NF-κB. The results in Figures 4D, 4E, and 5D, which focus on binding events that are most likely to be functionally relevant, also show co-occupancy by IκBζ, p50, and RelA. These results were somewhat surprising, as prior studies of IκBζ have proposed that it functions primarily by contributing a transactivation domain to p50 homodimers (Yamamoto et al. 2004; Trinh et al. 2008). However, other studies demonstrated that IκBζ interacts with both RelA and p50 in cell extracts, suggestive of an interaction with RelA:p50 heterodimers (Yamazaki et al. 2001; Totzke et al. 2006; Yamazaki et al. 2008; Kannan et al. 2011). Further uncertainty is provided by the fact that, despite the co-occupancy of putative functionally relevant sites by all three proteins, these sites often contain motifs capable of binding well to either p50 homodimers or RelA:p50 heterodimers (data not shown). For example, the putative *Il6* enhancer in Figure 5D contains the sequence, GGGGATCTCCCC, with 3 G:C-bps in each half-site, which has the capacity to bind diverse NF-κB dimers.

To further explore this issue, we performed RelA ChIP-seq in *Nfkb1^-/-^*BMDMs. The results revealed only a small subset of RelA binding events across the genome are dependent on p50 (Figure 6A). Many of these binding events correspond to genomic sites where IκBζ binding was found to be p50 dependent (Figure 6A, pink dots). To quantify this observation, RelA ChIP-seq peaks were divided into 20 bins on the basis of their p50 dependence. The percentage of peaks in each bin that exhibit p50-dependent binding of IκBζ was then calculated (Figure 6B). Remarkably, in the bin with RelA peaks showing the greatest p50-dependence, 27% of the peaks also exhibited p50-dependent IκBζ binding (Figure 6B). This finding further highlights the existence of a select subset of NF-κB genomic binding sites at which both RelA and IκBζ bind in a manner that is strongly dependent on the presence of p50. This contrasts with the vast majority of genomic sites where RelA and IκBζ binding are p50 independent. Although we have been unable to determine the key features that distinguish these two subsets of sites (other than the lower prevalence of consensus NF-κB motifs at sites where IκBζ binding is p50 independent), the results further support the possibility that IκBζ can bind DNA in association with RelA:p50 heterodimers.

**Figure 6.**
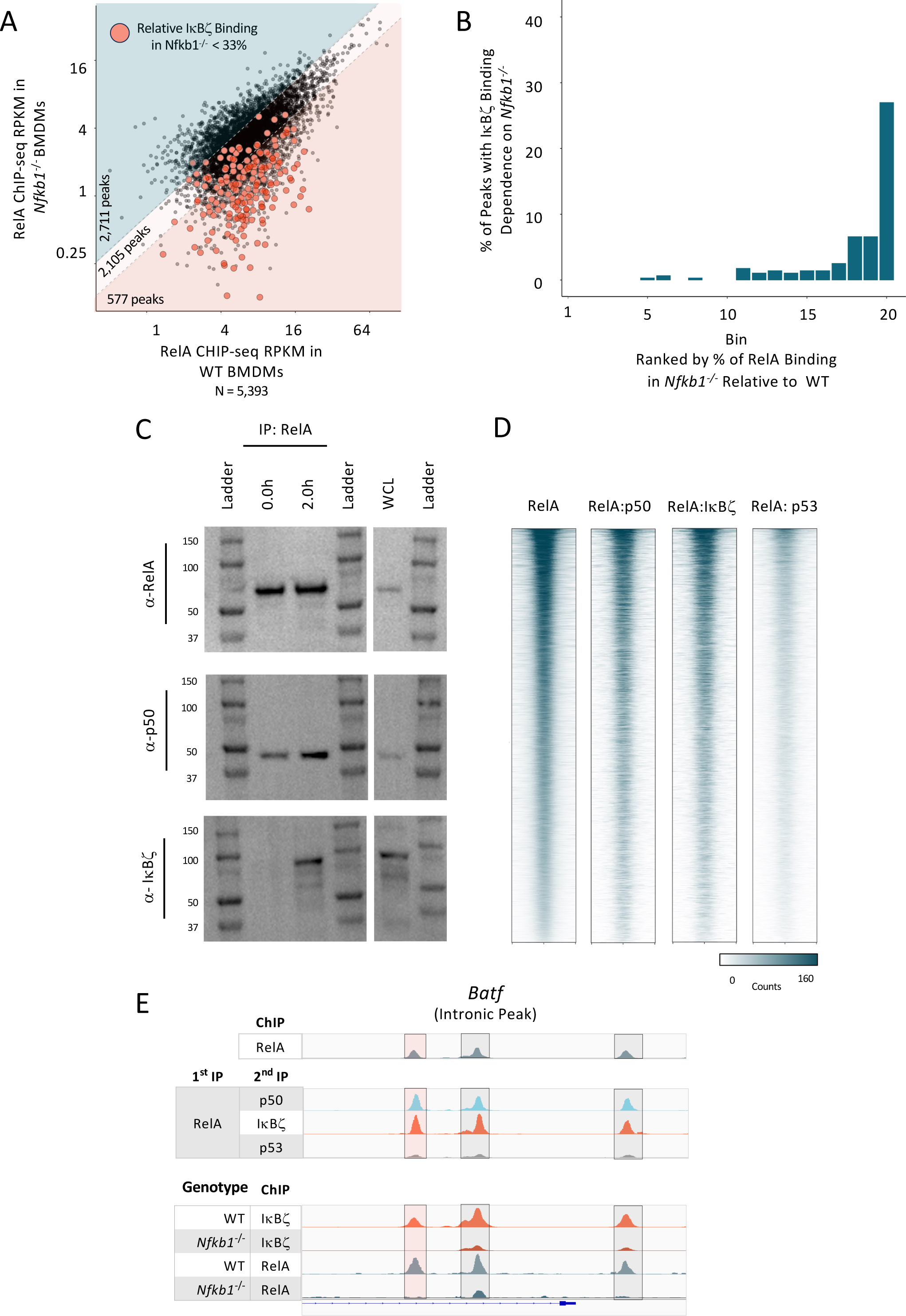
Possible Relevance of RelA in IκBζ-Dependent Transcription. (A) For all RelA ChIP-seq peaks (PS > 19 and RPKM >3 in 2/3 replicates) in WT macrophages stimulated with lipid A for 2 h, the average RPKM in WT (x-axis) versus *Nfkb1*^-/-^ (y-axis) macrophages is plotted. The pink background highlights 577 RelA peaks (10.7% of total) that exhibit the smallest *Nfkb1*^-/-^RPKM/WT RPKM ratios. Dots colored red indicate peaks that also display small *Nfkb1*^-/-^RPKM/WT RPKM ratios for IκBζ binding (<0.33, see Figure 5A). (B) Quantification of the results in panel A is shown. All RelA peaks are placed in 20 bins based on relative dependence on *Nfkb1*. In each bin, the percentage of peaks that annotate to locations that also exhibit *Nfkb1*-dependent IκBζ binding are displayed, showing that *Nfkb1*-dependent RelA binding frequently coincides with *Nfkb1*-dependent IκBζ binding. (C) Nuclear extract from BMDMs stimulated with lipid A for 0 or 2 hr were subject to immunoprecipitation using RelA antibodies. Immunoblot analysis was performed with the immunprecipitated material using anti-RelA, anti-p50, and anti-IκBζ antibodies. Whole-cell lysates (WCL) were examined by immunoblot as a control. (D) Heatmaps are shown for sequential ChIP-seq peaks. The first heatmap shows rank-ordered peaks obtained by RelA ChIP-seq. Heatmaps 2-4 show the same rank-ordered peaks after sequential ChIP-seq, with the first immunoprecipitation performed with RelA antibodies and the second immunoprecipitation performed with antibodies directed against p50 (heatmap 2), IκBζ (heatmap 3) or p53 (heatmap 4, as a negative control). (E) Browser tracks are shown for three sites in the *Batf* locus that exhibit *Nfkb1*-dependent binding of both RelA and IκBζ (The location highlighted in pink shows the greatest *Nfkb1*-dependence.) Track 1 shows RelA ChIP-seq peaks obtained in parallel with the sequential ChIP-seq results. Tracks 2-4 show the sequential ChIP-seq results using RelA antibodies for the first precipitation and p50 (track 2), IκBζ (track 3), and p53 (track 4) antibodies for the second precipitation. Tracks 5-8 show IκBζ or RelA ChIP-seq peaks from WT and *Nfkb1*^-/-^ BMDMs, as indicated.

The NF-κB dimer with which IκBζ associates at its functionally relevant sites in vivo is difficult to determine, largely because different protein complexes may be associated with a given genomic site at different alleles within a single cell or cell population. To address this challenge, we first performed co-immunoprecipitation experiments to examine IκBζ’s association with RelA in lysates from a J2-transformed BMDM line. Following immunoprecipitation with RelA antibodies (Figure 6C), immunoblots revealed an association with p50 in nuclear extracts from both unstimulated and stimulated cells, and an association with IκBζ only in nuclear extracts from stimulated cells, consistent with knowledge that IκBζ is poorly expressed prior to stimulation. Thus, our results are consistent with the prior studies that showed IκBζ interactions with RelA (see above).

We also performed electrophoretic mobility shift assays (EMSA) with extracts containing overexpressed RelA, p50, and IκBζ, either expressed individually and combined or co-expressed (we were unable to detect IκBζ -containing EMSA complexes in the absence of overexpression). As observed previously (Yamazaki et al. 2001; Totzke et al. 2006), IκBζ strongly inhibited DNA binding by both p50 homodimer and RelA:p50 heterodimers (data not shown). The inhibition of DNA-binding provides further evidence that IκBζ can bind both RelA:p50 heterodimers and p50 homodimers in vitro. However, in vivo, IκBζ clearly co-occupies DNA with NF-κB dimers rather than inhibiting their binding. One possible reason for the in vitro inhibition is that IκBζ may require processing or a post-translational modification for co-binding DNA with NF-κB dimers, with the processing event or modification absent in extracts containing overexpressed protein. Posttranslational modifications of IκBζ have been described (Grondona et al. 2020), but their functions remain poorly understood.

Finally, to further examine which NF-κB dimers can support IκBζ’s association with DNA in vivo, we employed sequential-ChIP-seq experiments. For these experiments, we first performed ChIP with RelA antibodies. We then used 10mM DTT to dissociate the immunoprecipitated chromatin from the antibody-bead complex, followed by a second immunoprecipitation with antibodies directed against either p50, IκBζ, or p53 (as a negative control). The results reveal that the use of p50 or IκBζ antibodies for the second immunoprecipitation resulted in comparably high efficiencies of immunoprecipitation of chromatin fragments previously immunoprecipitated with RelA antibodies, whereas p53 antibodies were much less efficient (Figure 6D). Browser tracks for three representative peaks at the *Batf* locus show the efficiency of secondary immunoprecipitation by the p50 and IκBζ antibodies in comparison to p53 antibodies (Figure 6E, tracks 2-4.). Notably, all three of these peaks exhibit substantial p50-dependent binding of both IκBζ and RelA (Figure 6E, tracks 5-8). These results therefore support the hypothesis that IκBζ has the potential to bind DNA in association with RelA:p50 heterodimers.

### Role of IκBζ in the Differential Responses to TLR4 and TNFR Signaling

As described above, a small set of key inflammatory and immunoregulatory genes exhibits strong p50/IκBζ transcriptional co-dependence. Other key genes, including *Il12b*, exhibit strong dependence on one or both factors in lipid A-stimulated macrophages. To explore the potential biological significance of the p50/IκBζ regulatory pathway, we searched for settings in which IκBζ might contribute to differential gene transcription. This effort led to a focus on BMDMs stimulated with lipid A versus TNF. Although *Nfkbiz* is a primary response gene that is potently activated by lipid A, previous studies have shown that the *Nfkbiz* gene is poorly expressed in TNF-stimulated macrophages (Yamazaki et al. 2001, 2004).

To extend the characterization of *Nfkbiz* differential expression, we performed nascent transcript RNA-seq and mRNA-seq with TNF-stimulated BMDMs. A comparative analysis reveals a surprisingly large magnitude of *Nfkbiz* differential expression between TNF and lipid A stimulation. *Nfkbiz* nascent transcripts and mRNA are 18-fold and 62-fold more abundant, respectively, after lipid A stimulation than after TNF stimulation at the 1-h time point (Figure 7A). The greater magnitude of the mRNA difference is likely due to a previously described instability of the *Nfkbiz* mRNA following TNF stimulation (Yamazaki et al. 2004), suggesting the existence of a mechanism to ensure that IκBζ levels remain extremely low following TNF stimulation.

**Figure 7.**
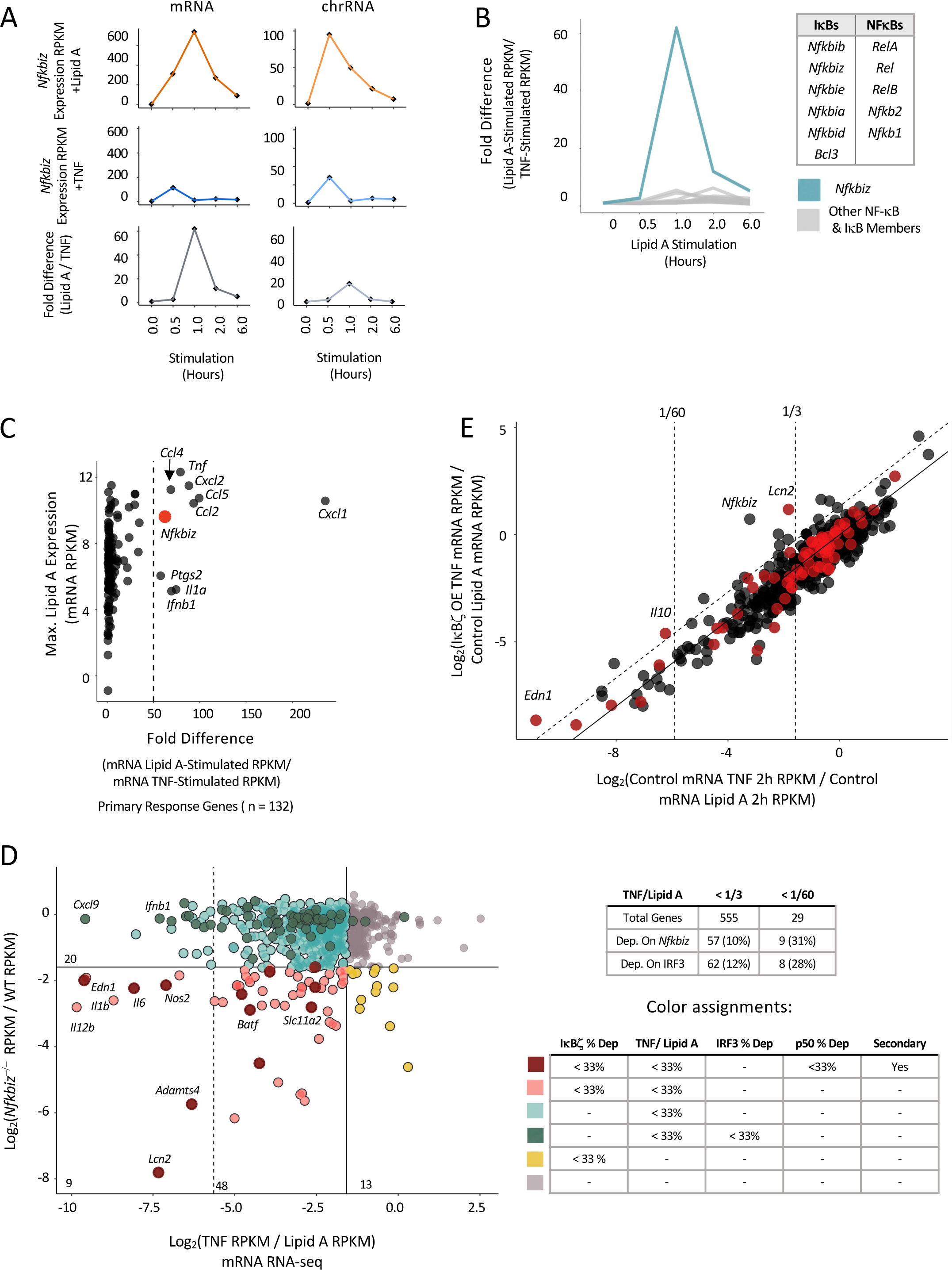
Contribution of IκBζ to Differential BMDM Responses to Lipid A and TNFα. (A) The line graphs show *Nfkbiz* transcript levels (RPKM) in stimulated BMDMs at each of five stimulation time points (0, 0.5, 1, 2, and 6 hr) from mRNA-seq (left) and nascent chromatin-associated transcript RNA-seq (right). Values are shown for lipid A (top) and TNF (middle) stimulation, with the lipid A/TNF ratio (fold difference) at each time point at the bottom. (B) The fold differences (ratios) between lipid A- and TNF-stimulated RPKM are shown at each time point for all five NF-κB and six IκB family members. *Nfkbiz* is highlighted in blue. (C) All 132 strongly induced (FC > 10) primary response genes (Tong et al. 2016) are plotted based on maximum mRNA transcript levels (RPKM) in lipid A-stimulated macrophages (y-axis) versus the fold-difference (ratio) in mRNA transcript levels between lipid A- and TNF-stimulated BMDMs (x-axis). The dashed line corresponds to a 50-fold difference*. Nfkbiz* is among only nine primary response genes that differ more than 50-fold. (D) A scatter plot shows that a large fraction of genes exhibiting the greatest differential transcription between lipid A- and TNF-stimulated BMDMs also exhibit strong *Nfkbiz*-dependent transcription. All 802 expressed and inducible genes (lipid A induction >3; WT RPKM >3) are plotted. The y-axis shows the ratio (log2) of chromatin-associated transcript levels (RPKM) in lipid A-stimulated *Nfkbiz*^-/-^ BMDMs versus lipid A-stimulated WT BMDMs. The x-axis shows the ratio of mRNA levels (RPKM) from TNF-versus lipid A-stimulated BMDMs. Vertical dashed lines represent TNF versus lipid A differential transcript ratios of 1/3 (right) and 1/60 (left). The horizontal dashed line represents an *Nfkbiz*^-/-^ /WT ratio of 1/3. Genes that exhibit both *Nfkbiz*-dependence and also display differential TNF versus lipid A transcript levels are in pink. Genes that exhibit *Nfkbiz* and *Nfkb1*-dependent, display differential TNF versus lipid A transcript levels, and are secondary response genes, are in dark red. Genes that are only *Nfkbiz*-dependent or that only display differential TNF versus lipid A transcript levels are in gold and light blue, respectively (see detailed color assignments at the bottom right). Note that 9 of 29 genes (31%) that exhibit the greatest differential TNF versus lipid A transcript levels (to the left of the left-most vertical dashed line) exhibit *Nfkbiz* dependence. At the right, the table shows the numbers of genes that exhibit *Nfkbiz*-dependence and IRF3-dependence among those genes that exhibit weak (<1/3) or strong (<1/60) TNF versus lipid A differential expression. (E) A scatterplot shows the impact of IκBζ overexpression on gene transcription in TNF-stimulated macrophages (a J2-transformed macrophage line). On the y-axis, for all lipid A-inducible genes (induction >3-fold; WT RPKM >3), ratios of mRNA levels (RPKM) in TNF-stimulated macrophages transduced with an *NFkbiz*-expressing retrovirus versus lipid A-stimulated untransduced macrophages are plotted (2-hr stimulation time point). On the x-axis, ratios of mRNA levels in TNF-stimulated macrophages versus lipid A-stimulated macrophages (2-hr time points) are plotted. Vertical dashed lines correspond to TNF versus lipid A differential transcript levels of 1/60 (left) and 1/3 (right). The diagonal solid line corresponds to unchanged gene expression in the presence of IκBζ overexpression. The diagonal dashed line corresponds to 2.5-fold higher transcript levels with TNF stimulation in the presence of IκBζ overexpression compared to untransfected cells stimulated with TNF. Genes that exhibit *Nfkbiz-*dependent expression (fold-change <33%; p-value <0.01) are highlighted in red. The *Lcn2* gene, which is impacted to the greatest extent by IκBζ overexpression, and the *Nfkbiz* gene, which is overexpressed from the retroviral vector, are noted.

The large magnitude of *Nfkbiz* differential expression is highly unusual. For example, *Nfkbiz* mRNA transcripts exhibit a differential expression magnitude that is much greater than that of all other NF-κB and IκB members (Figure 7B). Furthermore, among all 132 strongly induced (FC > 10) primary response genes activated by lipid A or TNF (Tong et al., 2016), *Nfkbiz* ranks ninth, with the others primarily corresponding to chemokines and cytokines rather than transcriptional regulators (Figure 8C). These results suggest that mechanisms have emerged to ensure that the abundance of IκBζ remains extremely low following TNF stimulation.

Consistent with the strongly differential expression of *Nfkbiz*, all of the p50/IκBζ co-dependent secondary response genes, including *Il6*, *Lcn2*, *Adamts4*, *Nos2*, and others (Figure 7D, left, maroon), and 81% of the IκBζ-dependent genes, including *Il12b* (Figure 7D, left, pink), exhibit strong differential nascent transcript levels (< 33% TNF/lipid A RPKM ratio). The 3 co-dependent primary response genes, including *Il1b*, also exhibit strong differential transcript levels (Figure 7D, left pink). In fact, among the genes showing the strongest differential expression (TNF/ lipid A percent expression < 2%; Figure 7D, genes to the left of the dashed vertical line), 31% are among the small group of p50/IκBζ or IκBζ targets.

To estimate the relative contributions of IRF3 and IκBζ to the differential responses of BMDMs to lipid A versus TNF, we used nascent transcript RNA-seq data sets from *Ifnar^-/-^* and *Irf3^-/-^* BMDMs to determine the number of genes that exhibit strong (< 33% relative to WT) dependence on IRF3 or its target, *Ifnb1*. This analysis revealed that 12% of lipid A vs. TNF differentially expressed genes exhibit IRF3/IFNAR-dependence (Figure 7D, top right). In comparison, 10% of lipid A vs. TNF differentially expressed genes exhibit IκBζ -dependence (Figure 7D, top right). Focusing more closely on the 29 genes that exhibit the strongest lipid A/TNF differential expression (nascent transcript level in response to TNF < 2% of the level in response to lipid A), we found that IRF3/IFNAR-dependence can account for the differential expression of 8 (28%) of these genes, with 11 (31%) exhibiting IκBζ -dependence (Figure 7D).

Finally, to determine whether an increased abundance of IκBζ is sufficient to rescue transcript levels for IκBζ-dependent genes in response to TNF stimulation, we overexpressed IκBζ in BMDMs by transduction of an IκBζ-expressing retrovirus, followed by stimulation with lipid A or TNF. In the presence of overexpressed IκBζ, the induction magnitude of one IκBζ-dependent gene, *Lcn2*, was comparable after TNF stimulation to the induction magnitude observed after lipid A stimulation, showing efficient rescue (Figure 7E). However, the differential induction of other p50/IκBζ co-dependent and IκBζ-dependent genes was not impacted by IκBζ overexpression (Figure 7E). One possible reason for the limited impact of IκBζ overexpression is that IκBζ activity may require complex regulatory mechanisms, such as posttranslational modifications (Grondona et al. 2020), that cannot be achieved with the overexpressed protein. The existence of such mechanisms is supported by the inability of recombinant IκBζ to co-occupy DNA with NF-κB dimers in EMSA experiments (see above).

## DISCUSSION

We performed a genomics-centric analysis of selective transcriptional regulation in stimulated BMDMs, with a focus on the NF-κB p50 protein and the nuclear IκBζ protein, IκBζ. The study began with the identification of a small group of key inflammatory and immunoregulatory genes that exhibited a strong dependence on p50 in nascent transcript RNA-seq experiments performed with lipid A-stimulated BMDMs. Strong overlap between p50-dependent and IκBζ-dependent genes revealed that a defined p50/IκBζ pathway makes a major contribution to the regulation of this key set of genes. Although p50, RelA, and IκBζ occupy thousands of genomic sites, a potentially defining characteristic of functionally meaningful binding events was the p50-dependence of IκBζ binding at a small subset of sites. Similar temporal kinetics of IκBζ binding and H3K27ac deposition also appeared to distinguish functional from non-functional IκBζ interactions. Sequential ChIP-seq and biochemical results provided strong support for the possibility that IκBζ can functionally interact with dimers containing p50 and also RelA, possibly RelA:p50 heterodimers. Biologically, the p50/IκBζ pathway contributes to the selective regulation of key immunoregulatory genes, some with pleiotropic functions. Moreover, this pathway appears to make a major contribution to the differential responses of macrophages to the microbial product, lipid A, and the inflammatory cytokine, TNF.

The small number of p50-dependent genes in BMDMs and the strong overlap with IκBζ-dependent genes was unexpected, given that p50 also contributes to stimulus-induced transcription as a component of the abundant RelA:p50 and c-Rel:p50 heterodimers. Redundancy between p50 and its closely related paralog, p52, provides a partial explanation for the limited number of p50-dependent genes, as a larger number of lipid A-induced genes exhibited reduced transcription in an *Nfkb1^-/-^Nfkb2^-/-^* macrophage line. However, the induction of most lipid A-induced genes was retained in these cells, suggesting that NF-κB dimers lacking p50 and p52 are sufficient for the induction of many genes. Importantly, p52 and other NF-κB family members are unable to compensate for the loss of p50 at a large fraction of IκBζ-dependent genes, showing an unusually close relationship between p50 and IκBζ.

Prior studies have shown that IκBζ binds p50 (Yamazaki et al. 2001; Yamamoto et al. 2004; Trinh et al. 2008; Kohda et al. 2014), but the fact that IκBζ dependence characterizes a high percentage of genes exhibiting a non-redundant dependence on p50 was not previously known. A subset of prior studies of IκBζ have suggested that its function involves association with p50 homodimers (Yamamoto et al. 2004; Trinh et al. 2008), with IκBζ providing an activation domain to a homodimer that otherwise would lack such a domain. However, our sequential ChIP-seq and biochemical results are more consistent with other prior studies that revealed in vitro interactions between IκBζ and both p50 and RelA (Yamazaki et al. 2001; Totzke et al. 2006; Yamazaki et al. 2008; Kannan et al. 2011). If RelA:p50 heterodimers carry out functional interactions with IκBζ, a mechanistic understanding of IκBζ’s role would require further exploration because RelA:p50 heterodimers possess RelA’s transactivation domain. According to combinatorial principles of gene regulation, the transcriptional activation of most if not all genes is thought to require multiple transcription factors, many with transactivation domains, and little is currently known about the mechanisms underlying the requirement for multiple transactivation domains for transcriptional activation. Thus, mechanistically, it is not unreasonable to envision a critical requirement for IκBζ for transcriptional activation when associated with a RelA:p50 heterodimer.

In addition to the need for further studies of IκBζ’s contribution to the activation of its defined target genes, studies are needed to understand why IκBζ’s interactions at the vast majority of its genomic binding sites have no apparent functional consequences. One finding that may be relevant to this issue is that most IκBζ genomic interactions are not p50-dependent. This finding suggests that IκBζ and its associated NF-κB dimer may adopt a specific conformation at a limited subset of sites that is critical for its transcriptional activation function.

Another unanswered question is why recombinant IκBζ appears to be incapable of binding DNA-associated p50 homodimers or RelA:p50 heterodimers in EMSA experiments, despite compelling evidence that it can associate with DNA-bound NF-κB dimers in vivo. This finding, observed by us and others (Yamazaki et al. 2001; Totzke et al. 2006), suggests that IκBζ may require a post-translational modification, a processing event, or the presence of another protein to allow association with DNA-bound NF-κB, raising the possibility of another layer of regulation of the p50/IκBζ pathway.

Although our data reveal strong overlap between p50-dependent and IκBζ-dependent genes, a substantial number of genes exhibit strong dependence on only one of these two proteins. A subset of these genes exhibits substantial dependence on both proteins but were not classified as co-dependent because they did not meet the stringent thresholds used in our study to define dependence. Nevertheless, some genes clearly exhibit strong dependence on one protein, with no impact of the other factor in loss-of-function experiments. A subset of IκBζ-dependent/p50-independent genes can be attributed to redundancy between p50 and p52, but it is not known whether other IκBζ-dependent/p50-independent genes collaborate with other NF-κB dimers or perhaps contribute to gene regulation in concert with a different transcription factor family. p50-dependent/IκBζ -independent transcription is likely to reflect a requirement at a small set of genes for a p50 dimeric species that does not require IκBζ for its function.

The key immunoregulatory genes that exhibit p50/IκBζ co-dependence, including *Il6* and *Il1b*, are potently activated in a large number of cell types in response to diverse stimuli. We cannot exclude the possibility that p50 and IκBζ are universally required for the activation of these genes, but many genes are regulated by different factors in different biological settings. Although a broad range of abnormalities have been reported in *Nfkb1^-/-^* and *Nfkb2^-/-^*mice (Hayden and Ghosh 2013), even more abnormalities would likely be observed if p50 and IκBζ were universally required for the transcription of the target genes identified in this study.

Finally, the finding that the *Nfkbiz* gene and most IκBζ target genes stand out due to their very large magnitude of differential expression between lipid A- and TNF-stimulated cells highlights the potential biological importance of the differential expression of these genes. The unusually low expression of *Nfkbiz* in TNF-stimulated macrophages appears to be due to the combined influence of a transcriptional mechanism that is unique among primary response genes and an mRNA stability mechanism that also appears unusual or unique (Yamazaki et al. 2004). The well-documented importance of many IκBζ target genes for anti-microbial responses provides justification for their potent activation by lipid A, but the reason transcriptional and post-transcriptional mechanisms evolved to ensure that these genes remain largely silent in response to TNF signaling is less obvious. Nevertheless, the large percentage of IκBζ-dependent genes that exhibit differential expression in response to lipid A versus TNF, and the unusually large magnitudes of their differential expression in comparison to the magnitudes observed with IRF3-dependent genes, suggests that IκBζ and the p50/IκBζ pathway are major contributors to differential lipid A/TNF gene induction in macrophages.

### EXPERIMENTAL PROCEDURES

#### Mice

The *Nfkb1*^-/-^ and *Nfkb2*^-/-^ mice were a gift from Alexander Hoffmann at UCLA. The *Nfkbiz*^-/-^ (Yamamoto et al. 2004), *Bcl3^-/-^* (Franzoso et al. 1997B), and *Nfkbid^-/-^* (Touma et al. 2007) mice, all bred onto a C57BL/6 background, were kindly provided by Giorgio Trinchieri (NIH), Ulrich Siebenlist (NIH) and Ingo Schmitz (Ruhr-University Bochum, Germany), respectively.

#### BMDM Isolation, Differentiation, and Stimulation

Bone marrow was extracted from male mice aged 8-12 weeks and differentiated into BMDMs as described (Ramirez-Carrozzi et al. 2009; Bhatt et al. 2012). In brief, following mouse euthanasia, the tibia, femur, and hip bones were removed. Bone marrow was flushed with a 25-gauge needle filled with 10 mL of PBS. Bone marrow was passed through a 70-uM filter, pelleted for 10 mins at 1300 rpm, and resuspended in 1 mL per mouse of RBS Lysis Buffer (Sigma, #R7757). Following the removal of red blood cells, the cells were pelleted and resuspended in media containing 20% FBS, 10% CMG, 1X Pen-Strep (Gibco, #15140-122), 1× L-Glutamine (Gibco, #25030-081), and 0.5 mM sodium pyruvate (Gibco, #11360-070) in Dulbecco’s Modified 44 Eagle Medium. 10 × 10^6^ cells were plated in 20 mLs of media on a 15 cm plate. Media was replaced on day 4. Macrophages were treated on day 6 with 100 ng/mL Lipid A (Sigma, L6895) or 10 ng/mL TNF (Bio-Techne, 410-MT). For MAPK inhibitor-treated samples, macrophages were pretreated with 1 uM ERK-inhibitor (PD 0325901) for 1 hr prior to stimulation.

#### RNA-seq

For chromatin-associated RNA-seq, approximately 15 million BMDMs per sample were stimulated and harvested on day 6 of differentiation. Chromatin-associated RNA and mRNA samples were prepared as described (Bhatt et al. 2012; Tong et al. 2016). In brief, for chromatin-associated RNA, cellular fractionation was used to isolate chromatin, which was resuspended in TRIzol reagent (ThermoFisher, #15596018). RNA was extracted with Qiagen RNeasy plus mini kit (Qiagen #74136). During purification, columns were treated with DNAse I to eliminate genomic DNA. For library preparation, 60 ng or 500 ng of RNA was used for chromatin-associated RNA or mRNA, respectively, to generate strand-specific libraries with the TruSeq stranded RNA Kit (Illumina, #RS-122-2001). The library preparation protocol was adapted for chromatin-associated RNA-seq (Bhatt et al. 2012). Libraries were sequenced using single-end (50 bps) Illumina Hi-Seq 2000 or 3000. Reads were aligned to the NCBI37/mm9 genome using Hisat2 (Kim et al. 2015; Kim et al. 2019). Following alignment, SAMtools was used to compress, sort, and index files (Danecek et al. 2021). HOMER software allowed for further processing and visualization of the data on the UCSC genome browser (Heinz et al. 2010). Using IGV Tools we generated TDFs for data visualization on IGV (Robinson et al. 2011). SeqMonk was used to extract read counts from BAM files (Babraham Bioinformatics). Read counts were normalized to gene size (kbps) and depth of sample sequencing (per million reads) to generate RPKMs.

#### ChIP-Seq and Sequential ChIP-seq

For ChIP-seq experiments, approximately 30 million (45 million for sequential ChIP-seq) BMDMs were harvested on day 6 of differentiation (two 15-cm plates per sample). RelA (Cell Signaling, 8242S), p50 (Cell Signaling, 13586), IκBζ (4301779, Sigma), and H3K27ac (Active Motif 39133) antibodies were used for immunoprecipitation. For sequential ChIP-seq, after the first immunoprecipitation, the protein-DNA complex was eluted from the antibody-bead complex with 10 mM DTT at 37°C for 30 min. The eluent was then diluted 20X, the second antibody was added, and the ChIP-seq protocol was continued as described (Barish et al. 2010). Reads were aligned to the NCBI37/mm9 genome using Hisat2 (Kim et al. 2015; Kim et al. 2019). Files were further compressed, sorted, and indexed using SAMTools (Danecek et al. 2021). IGV Tools was used to generate TDFs for data visualization on IGV (Robinson et al. 2011). Peaks were called with HOMER software using a false discovery rate of 0.01 (Heinz et al. 2010). BEDTools software allowed us to create a comprehensive list of all overlapping peaks between samples (Quinlan et al. 2010). We extracted raw read counts from BAM files with SeqMonk (Babraham Bioinformatics). Read counts were normalized to peak size (kbps) and depth of sample sequencing (per million reads) to generate RPKMs. Only reproducible peaks with the described criteria were maintained for downstream analysis.

#### ATAC-seq

ATAC-seq libraries were prepared with the Nextera Tn5 Transposase kit (Illumina) as described (Tong et al. 2016; Buenrostro et al., 2015). Reads were mapped to the NCBI37/mm9 genome using Hisat2 (Kim et al. 2015; Kim et al. 2019). SAMTools was used to compress, sort, index, and remove duplicates from samples. MACS2 was used to call peaks using a false discovery rate of 0.01 (Zhang et al. 2008). To create a complete list of accessible chromatin regions in all samples, we used BEDTools (Quinlan et al. 2010). We extracted raw read counts from BAM files with SeqMonk (Babraham Bioinformatics). Read counts were then normalized to peak size (kbps) and depth of sample sequencing (per million reads) to generate RPKMs.

#### CRISPR/Cas9 Mutagenesis

Synthetic guide RNAs (gRNAs) were designed using CRISPOR (http://crispor.tefor.net/) and Massachusetts Institute of Technology CRISPR Designer (http://crispr.mit.edu). Recombinant Cas9 (Synthego) in combination with gRNAs were electroporated into J2-transformed *Nfkb1*^-/-^ macrophages to delete exon 3 of *Nfkb2.* Single-cell clones were screened by genotyping and confirmed by immunoblot for p52 and p50 (Cell Signaling D9S3M and D4P4D) and by sequencing.

#### Motif Analysis

HOMER software was used to search for de novo motifs and the enrichment of position weight matrices for NF-κB from the Jaspar motif database (Heinz et al. 2010). The 200-bps surrounding the center of p50, RelA, or IκBζ ChIP-seq peaks were used in motif analyses.

#### IκBζ Retroviral Overexpression

The IκBζ expression construct was prepared in a pMSCV vector by VectorBuilder (https://en.vectorbuilder.com/). The construct was verified with DNA sequencing. Viral production was carried out by VectorBuilder and prepared to > 10^7^ TU/mL. Retroviral transductions were done as described (Sanjabi et al. 2005). In brief, bone marrow was collected from mice (described above) and plated in a 6-well dish at a density of 0.5 X 10^6^ per mL in CMG-conditioned media. On days 1-3, spin infections were performed. Cells were spun at 2500 rpm for 5 min, the supernatant was removed, media containing the virus, 8 μg/mL polybrene, and 10 μL/mL of 1M HEPES (pH 7.55) was incubated with cells for 1.5 hours at 2500 rpm at 4°C. After spin infections, virus-containing media was removed and CMG-conditioned media was added. On day 6, cells were stimulated and collected for either western blotting or RNA-seq.

## SUPPLEMENTAL FIGURE LEGENDS

**Supplemental Figure S1.**
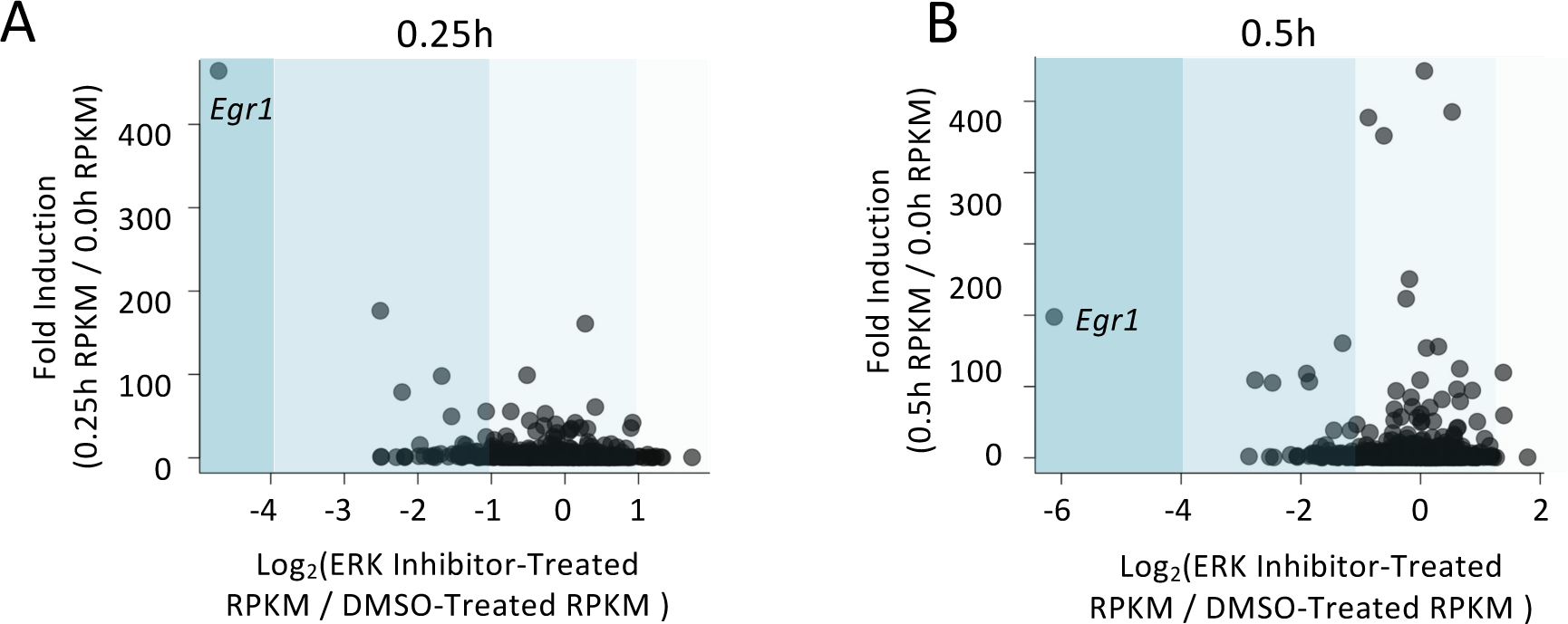
Uniquely Strong Dependence of *Egr1* Transcription on ERK Signaling. BMDMs were pretreated for 1 hr with ERK inhibitor, PD-0325901 (final concentration 5 μM dissolved in DMSO) or DMSO and subsequently stimulated with lipid A. mRNA-seq was then performed with cells stimulated with lipid A for 0.25 or 0.5 hr. The scatter plot shows, on the x-axis, the log2 ratio of RPKM in ERK-inhibitor treated cells versus control cells for the 132 strongly induced primary response genes (Tong et al. 2016). The y-axis shows the fold-induction of the 132 primary response gene transcripts at the relevant time point.

**Supplemental Figure S2.**
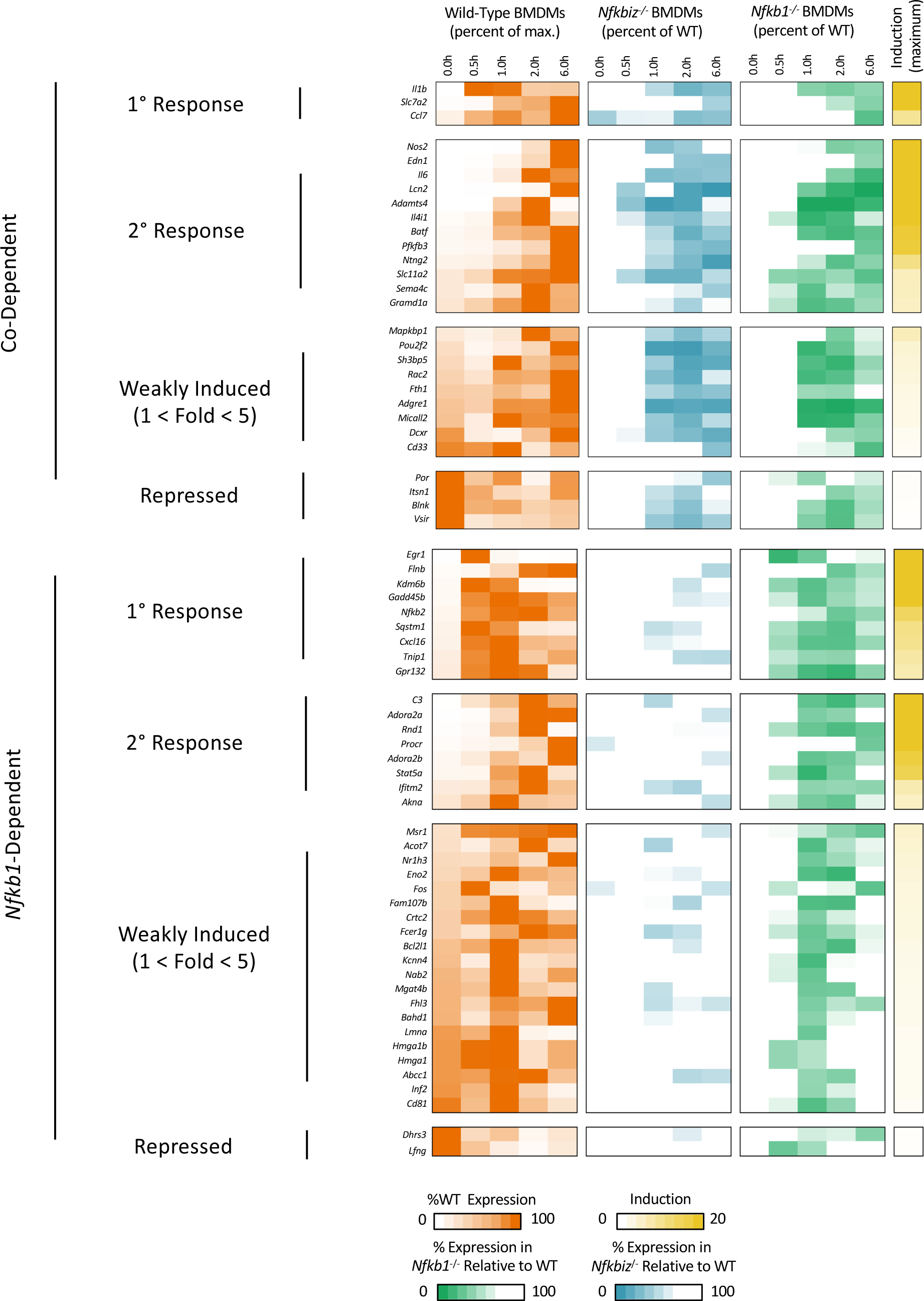
Lists of *Nfkb1/Nfkbiz*-Co-Dependent and *Nfkb1*-Dependent Genes in Lipid A-Stimulated Macrophages. Gene names and heatmaps of lipid A activation kinetics are shown (% of maximum RPKM) for the 28 *Nfkb1/Nfkbiz*-co-dependent and 39 *Nfkb1*-dependent genes. Dependence is defined as <33% chromatin-associated transcript levels at the 0.5-, 1-, 2-, or 6-hr time points in mutant cells in comparison to WT cells examined in parallel, in addition to a p-value <0.01 from two biological replicates. The genes are classified as primary response (those genes defined as primary response in Tong et al. 2016, plus additional genes induced >5-fold at any time point, with RPKM >3 at any time point, and with >33% chromatin-associated transcript levels in the presence of CHX at any time point), secondary response (those genes defined as secondary response in Tong et al. 2016 plus additional genes induced >5-fold at any time point, with RPKM >3 at any time point, and with <33% transcript level in the presence of CHX at any time point), weakly induced (those genes that not defined as primary or secondary response, but with RPKM >3 at any time point and with maximum induction in WT BMDMs of 1-5-fold), or repressed (those genes with RPKM >3 at any time point and with repression rather than induction in WT BMDMs). Heatmaps show the percent transcript level for each gene relative to the maximum transcript level observed for that gene at any time point in WT cells. The final heatmap column shows the maximum fold induction for each gene relative to the fold induction for all other genes.

**Supplemental Figure S3.**
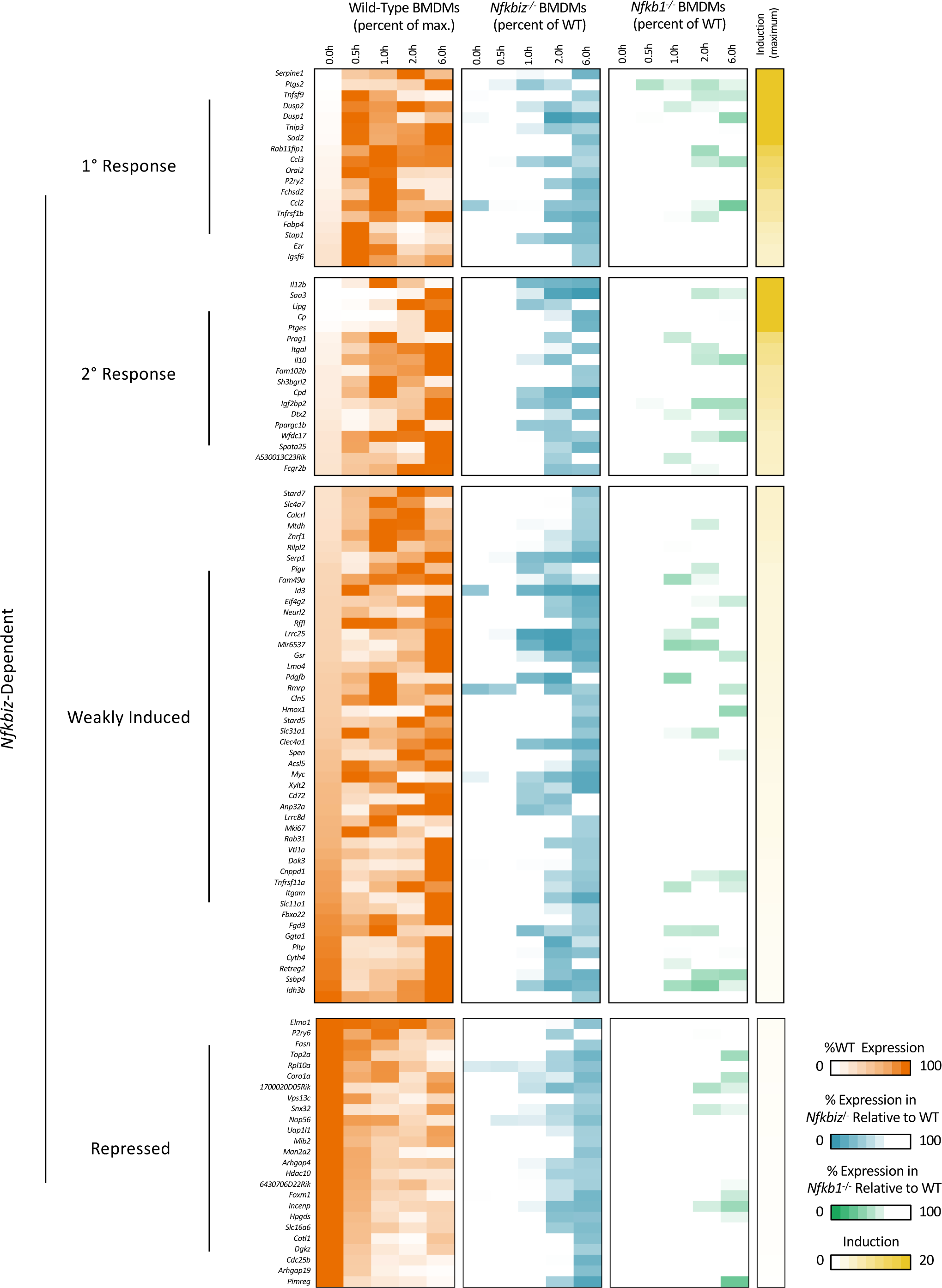
A List of *Nfkbiz*-Dependent Genes in Lipid A-Stimulated Macrophages. Gene names and heatmaps of lipid A activation kinetics are shown for the 108 genes with only *Nfkiz*-dependence. Definitions of dependence, classification criteria, and presentation details are the same as in Supplemental Figure S2.

**Supplemental Figure S4.**
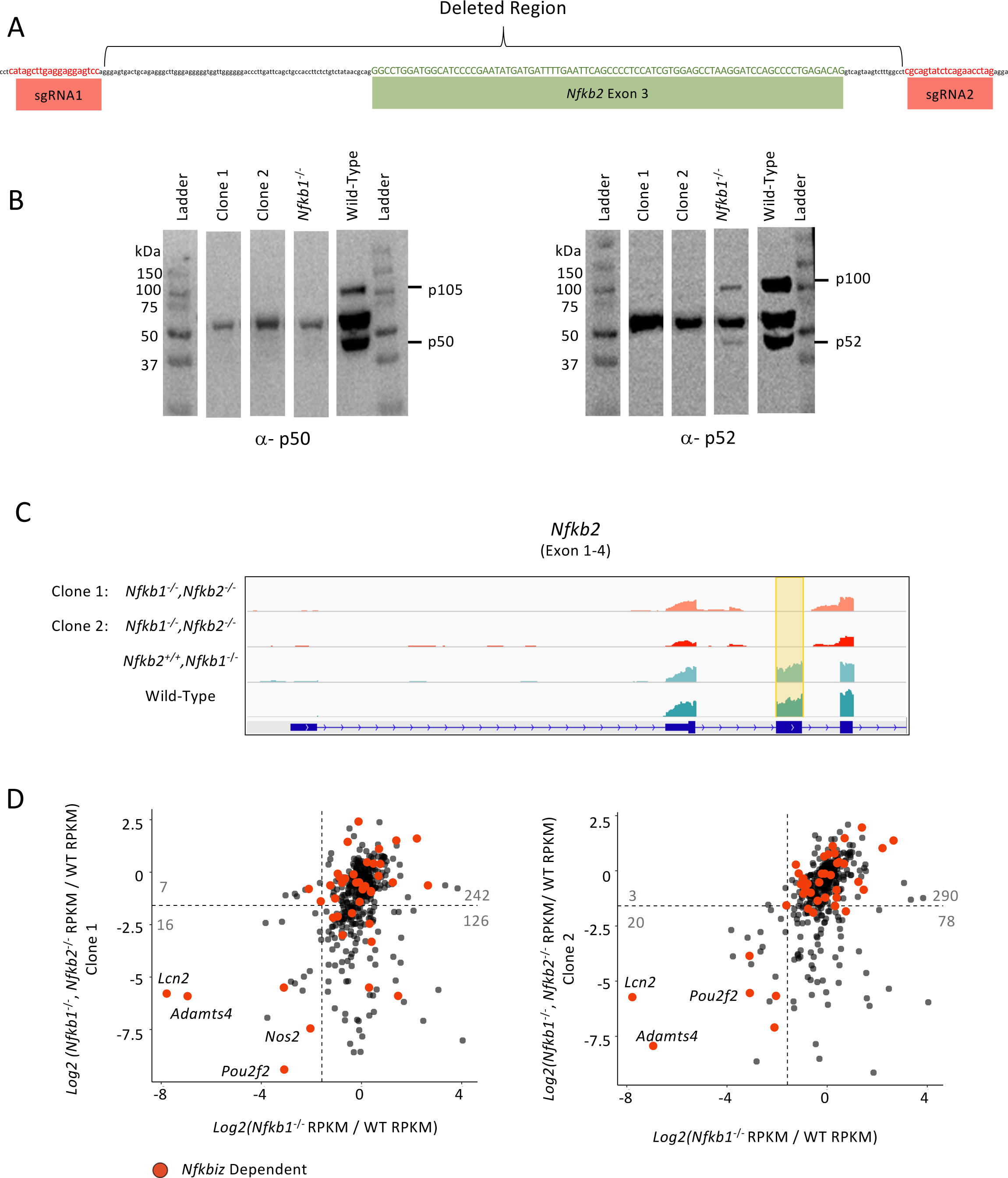
Generation and Analysis of Nfkb1^-/-^Nfkb2^-/-^ J2-Transformed Macrophage Lines. BMDMs from *Nfkb1^-/-^* mice were transformed with the J2 retrovirus. A homozygous deletion was then introduced into the *Nfkb2* gene by CRISPR/Cas9 mutagenesis, following the protocol described in Feng et al. 2024. Two clonal *Nfkb1^-/-^Nfkb2^-/-^* lines (clones 1 and 2) were selected and analyzed. (A) The sequence of the region surrounding *Nfkb2* exon 3 is shown, along with the two single guide RNAs (sgRNAs) used for mutagenesis. (B) Immunoblot analyses were used to confirm the absence of both the p50 (left) and p52 (right) proteins in *Nfkb1^-/-^Nfkb2^-/-^* clones 1 and 2, with *Nfkb1^-/-^* cells and WT cells examined as controls. The p50 and p52 antibodies recognized the processed (p50 and p52) and precursor (p105 and p100) proteins translated from the two genes, as well as a non-specific protein. (C) The deletion of *Nfkb2* exon 3 was further confirmed by examination of mRNA-seq tracks from clones 1 and 2, with *Nfkb1^-/-^*cells and WT cells examined as controls. (D) Scatter plots compare the impact on mRNA levels of lipid A-induced genes in *Nfkb1^-/-^* and *Nfkb1^-/-^Nfkb2^-/-^* macrophage lines. mRNA-seq was performed with two independent *Nfkb1^-/-^Nfkb2^-/-^* lines, clone 1 (left) and clone 2 (right), the *Nfkb1^-/-^* parental line, and a WT J2-transformed line. The x-axes correspond to the *Nfkb1^-/-^* RPKM/WT RPKM ratio for each lipid A-induced gene (induction >5-fold), and the y-axes correspond to the *Nfkb1^-/-^Nfkb2^-/-^*RPKM/WT RPKM ratio. *Nfkbiz*-dependent genes are in red. Dashed lines correspond to ratios of 0.33.

